# Early Binding of Anti-Amyloid Antibodies to CAA Drives Complement Activation, Inflammation and ARIA in Mice

**DOI:** 10.64898/2026.03.04.709591

**Authors:** Praveen Bathini, Stephan Schilling, Jens-Ulrich Rahfeld, David M. Holtzman, Takaomi C Saido, Cynthia A. Lemere

**Author notes:** Correspondence to: Cynthia A. Lemere;, Praveen Bathini.

## Abstract

Anti-amyloid antibody treatment for Alzheimer’s disease is linked to Amyloid-Related Imaging Abnormalities (ARIA), including vasogenic edema (ARIA-E) and microhemorrhages (ARIA-H), especially in ApoE ε4/4 carriers. To investigate mechanisms underlying ARIA, we examined the binding and temporal vascular effects of immunization with 3D6, the precursor to the anti-amyloid antibody bapineuzumab, in two aged Alzheimer’s disease amyloid mouse models. Acutely, 3D6 bound to cerebral amyloid angiopathy (CAA), resulting in C1q binding and classical complement activation. Weekly short-term immunization over 7 weeks resulted in elevated CAA- and plaque-associated complement deposition, red blood cell extravasation and microhemorrhages, and was accompanied by significant transcriptomic changes in genes related to complement, inflammation, vascular dysfunction, and endothelial lipid responses. Longer-term dosing over 13-15 weeks further increased complement deposition and was associated with blood-brain barrier disruption, MMP-9 upregulation, and microhemorrhages, accompanied by reduced amyloid burden and modest CAA clearance. C3 levels correlated with microhemorrhage severity. Perivascular macrophages co-localized with complement-decorated CAA in 3D6-treated mice. These findings implicate complement activation as an early key driver of ARIA and suggest that therapeutic targeting of complement may reduce ARIA risk.

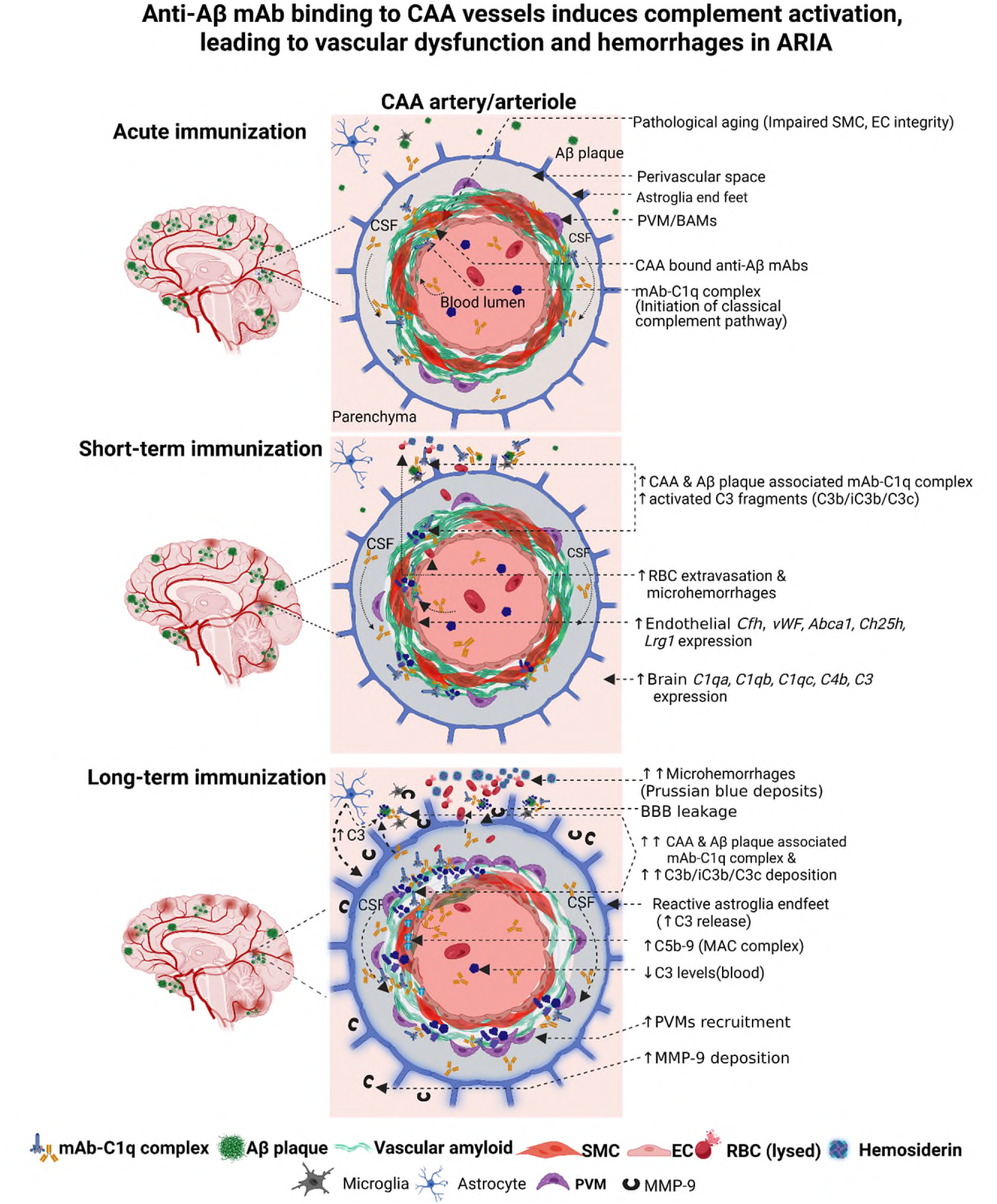

## Introduction

Alzheimer’s disease (AD) is a progressive neurodegenerative disorder characterized by extracellular amyloid-β (Aβ) plaques and intracellular hyperphosphorylated tau tangles that ultimately drive cognitive decline [1]. Because Aβ accumulation is an early event in AD pathogenesis, passive immunotherapy with anti-Aβ monoclonal antibodies (mAbs) has emerged as a strategy to slow disease progression [2]. Bapineuzumab, the first anti-Aβ mAb to enter clinical trials, revealed on MRI a spectrum of amyloid-related imaging abnormalities (ARIA) comprising vasogenic edema or effusion (ARIA-E) and microhemorrhages and superficial siderosis (ARIA-H), linked to vascular leakage and red blood cell extravasation [3,4]. Subsequent second-generation mAbs, aducanumab (Aduhelm), lecanemab (Leqembi), and donanemab (Kisunla) demonstrated robust amyloid clearance and modest slowing of cognitive decline, but all were associated with ARIA-E and ARIA-H in subsets of treated patients [5,6], raising safety concerns.

ARIA remains incompletely understood and there is an urgent need to understand the mechanisms underlying the pathology [5–7]. ARIA events occur most frequently during early treatment phases [8–10], and their risk is heightened by high antibody doses, ApoE ε4 allele carriage, and pre-existing cerebral amyloid angiopathy (CAA) [11,12]. CAA, defined by amyloid deposition within the walls of cerebral vessels, is common in AD [13] and is exacerbated by the ApoE ε4 genotype, which increases vascular lesion burden [14,15] and spontaneous intracerebral hemorrhage risk [16]. These observations suggest a synergistic interaction between CAA load, ApoE4-associated vascular fragility, and susceptibility to ARIA. Indeed, CAA may be necessary for ARIA to occur and may amplify its pathology [17]. Clinically, ApoE ε4 homozygous patients showed higher rates of ARIA-E (lecanemab: 32.6% in ε4/ε4 vs 10.9% in ε4/–; donanemab: 41.1% vs 23.8%) and ARIA-H (lecanemab: 39% vs 14%; donanemab: 53.6% vs 30.8%) [8,18], with most events emerging between the 4th–8th infusion for lecanemab [19] or 1st–6th for donanemab [8]. Although often asymptomatic and mild, severe ARIA can be fatal. For example, a 79-year-old ε4/ε4 patient on lecanemab developed fatal β-amyloid-related arteritis after the third infusion, likely exacerbated by anticoagulation treatment [20]. In another ε4/ε4 AD patient treated with lecanemab, computed tomography before tissue plasminogen activator (t-PA) anti-coagulant administration showed hypodensities in the left temporal–parietal regions and a distal middle cerebral artery occlusion without hemorrhage consistent with acute ischemic stroke. However, subsequent t-PA treatment led to multifocal intracerebral hemorrhages and necrotizing vasculopathy, resulting in death [21].

Consistent with clinical findings, passive anti-amyloid immunization has been shown to induce microhemorrhages in multiple AD amyloid mouse models bearing CAA [22–24]. Notably, N-terminally-directed anti-amyloid antibodies that bound deposited Aβ exacerbated CAA-associated vascular damage compared with central-domain antibodies which only recognized Aβ monomers [23], underscoring the impact of whether the antibodies can recognize Aβ species present in Aβ deposits.

Several mechanisms have been postulated for ARIA. For example, anti-Aβ mAbs may promote rapid plaque disaggregation and redistribution of soluble Aβ into vessel walls, overwhelming perivascular drainage [3]. Or, they bind directly to vascular amyloid, compromising vessel wall integrity and causing fluid extravasation [3,25]. In addition, they may activate glial and perivascular macrophages, promoting vascular inflammation [26]. The pathological overlap between antibody-induced ARIA and CAA-related inflammation (CAA-ri), wherein CAA is associated with perivascular inflammation [11], suggests some shared mechanisms. In both ARIA and CAA-ri, complement activation emerges as a likely effector pathway. Complement proteins including C1q, C4, and activated C3 fragments localize to plaques [27] and CAA-affected vessels [28], while the terminal membrane attack complex (C5b-9) accumulates in CAA-laden arteries and arterioles, correlating with subcortical hemorrhage and cortical superficial siderosis [28]. The classical complement pathway is triggered when C1q binds to the Fc region of an antibody once it is bound to its antigen, forming the C1 complex with serine proteases C1r and C1s, which cleaves C4 and C2 to generate C3 convertase (C4b2b), which then cleaves C3 into anaphylatoxin C3a for immune cell recruitment and release of pro-inflammatory molecules, and opsonic C3b/iC3b for phagocytosis upon binding to its receptor CR3 on macrophages and microglia [29]. Downstream cleavage of C5 by C5 convertases (C4b2b3b, classical pathway and C3bBbC3b, alternative pathway) produces the anaphylatoxin C5a and C5b which binds C6, C7, C8 and multiple C9 molecules to form the terminal membrane attack complex C5b-9 which causes cell lysis [30,31].

Given the convergence of ARIA, CAA-ri, and complement activation, we hypothesized that early anti-Aβ antibody binding to CAA triggers the activation of the classical complement pathway, and that repeated exposure amplifies vascular inflammation and damage. To test this hypothesis, we passively immunized mice with recombinant 3D6, the murine IgG2a precursor to bapineuzumab, and examined antibody binding and complement activation in brain across different dosing paradigms, treatment durations, and amyloid mouse models of AD to delineate when and where complement engagement contributes to antibody-induced cerebrovascular injury. By dissecting this interplay between CAA, anti-amyloid antibodies and complement, we identify complement effectors as potential therapeutic targets for mitigating ARIA in AD.

## Results

### Anti-Aβ mAb 3D6 bound first to vascular amyloid and was associated with complement C1q deposition

CAA occurs mostly in arterioles in the leptomeninges and in large penetrating arterioles, predominantly in cerebellum and to a lesser degree in cortex, and in arterioles near the ventricles in the amyloid mouse models used in these studies. To assess antibody binding to amyloid and subsequent complement activation, both 3D6 and IgG2a isotype control mAbs were biotinylated and administered intraperitoneally (i.p.) as a single 25 mg/kg dose to 19-month-old *APP^NL-G-F^* mice, followed by euthanasia 24 h later (Fig. 1a). Streptavidin labeling indicated that 3D6-Biot preferentially bound vascular amyloid (arrowheads, Fig. 1b) in the penetrating cortical pial vessels. Parenchymal plaque labeling by 3D6 was absent at this stage (asterisks). Notably, 3D6 binding was accompanied by C1q deposition in pial vessels and large arterial vessels bearing amyloid near the ventricles (Fig. 1c,d).

**Fig. 1:**
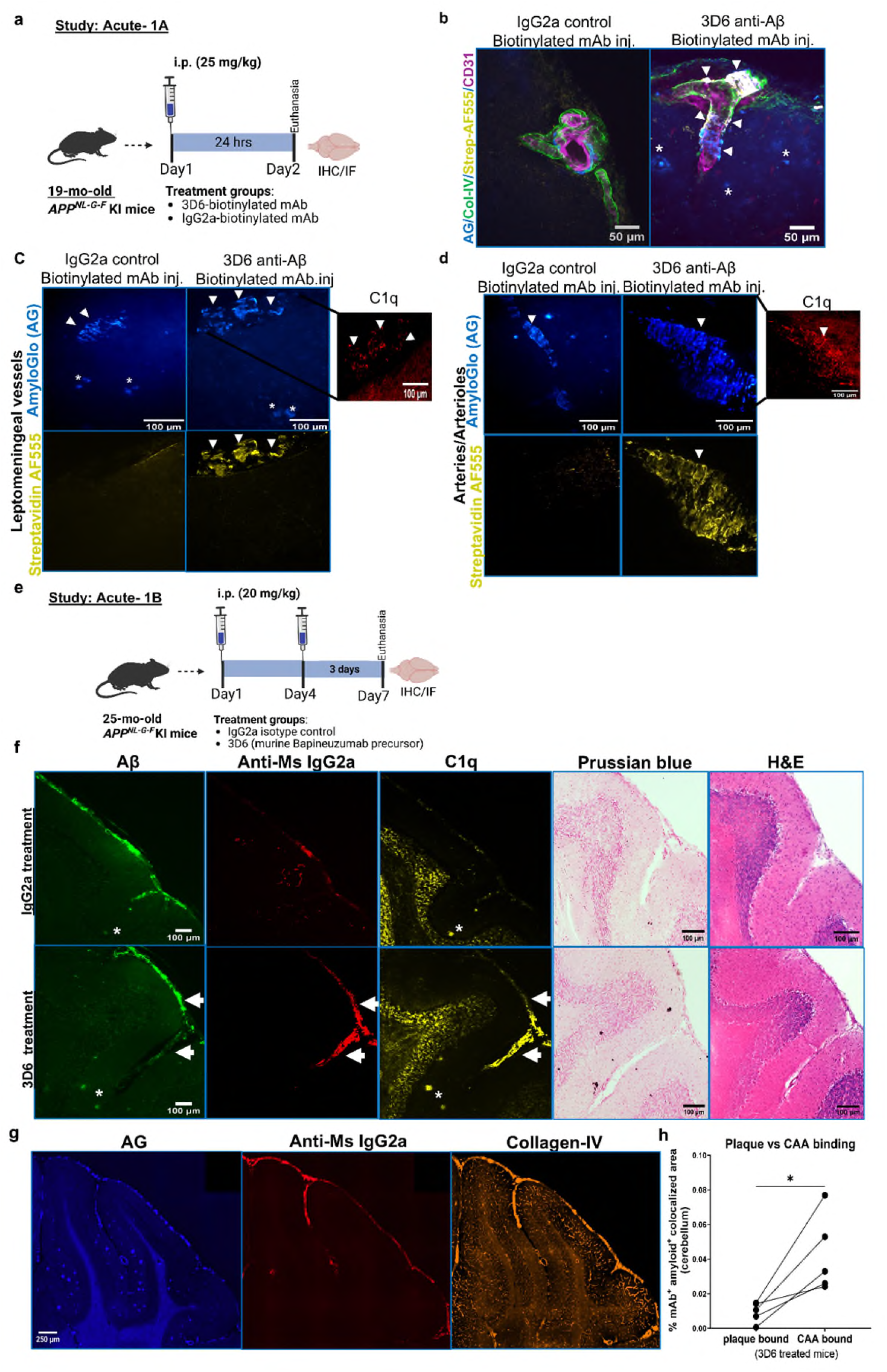
Acute anti-Aβ mAb passive immunization with 3D6, the murine bapineuzumab precursor, demonstrated early preferential antibody binding to CAA. **a,** Schematic of acute single-dose immunization in 19-month-old *APP^NL-G-F^* KI mice (n=3-4/group) with biotinylated 3D6 or IgG2a mAb (25 mg/kg, i.p.), followed by euthanasia at 24 h. **b,** Brain sections stained with Amylo-Glo (AG) for amyloid, Collagen-IV for blood vessels’ basement membrane, Streptavidin 555 for biotinylated 3D6 or IgG2a, and endothelial marker CD31 in IgG2a and 3D6-biotinylated mAb-injected mice; images taken from the leptomeningeal penetrating vessels along the cortical pial surface. Arrowheads point towards CAA-bound Streptavidin 555, indicative of biotinylated mAb binding; asterisks indicate plaques. **c–d**, Brain sections labeled with AG and Streptavidin-AF555 showing biotinylated mAb distribution in **c,** leptomeningeal vessels, and d, large parenchymal arterioles. Arrowheads indicate CAA-associated mAb binding and C1q deposition observed in 3D6-treated mice. **e,** Schematic of acute immunization in 25-month-old *APP^NL-G-F^* KI mice (n= 7-9/group) treated with anti-Aβ 3D6 IgG2a mAb or IgG2a isotype control. **f,** Brain sections stained for aggregated Aβ, anti-mouse IgG2a (to detect 3D6 and IgG2a), and C1q in IgG2a control- and 3D6-treated mice. Arrows indicate cerebellar meningeal vessel-associated labeling; asterisks mark parenchymal plaque labeling. Adjacent sections were stained with Prussian blue (counterstained with nuclear fast red) for microhemorrhages and H&E for RBC extravasation. **g,** Triple labeling with AG, anti-mouse IgG2a, and collagen-IV to distinguish plaque (AG⁺AntiMsIgG2a⁺ColIV⁻) and vascular (AG⁺AntiMsIgG2a⁺ColIV⁺) mAb binding in 3D6-treated brain sections. **h,** Quantification of % cerebellar plaque vs CAA labeling of mAb in a subset of 3D6 treated mice (n=5), *p < 0.05, paired t-test. Scale bar: b=50 µm; c,d,f =100 µm; g = 250 µm.

To further evaluate early antibody binding, complement activation, and ARIA-like changes, 25-month-old *APP^NL-G-F^* mice were administered with 20 mg/kg 3D6 or IgG2a on Day 1 and Day 4, followed by euthanasia on Day 7 (Fig. 1e). Immunolabeling for aggregated Aβ, anti-mouse IgG2a as a surrogate marker of the immunizing mAbs, and C1q revealed strong CAA, mAb and C1q colocalization in cerebellar meningeal sulcal vessels (arrows, Fig. 1f) in 3D6-treated mice, but not in IgG2a controls. Mab-binding to plaques was relatively low at this timepoint, although C1q deposition in plaques (asterisks) was seen in both groups, consistent with previous reports of C1q colocalization with plaques in amyloid mice and human AD brains [28,32]. No red blood cell (RBC) extravasation or microhemorrhages were observed by hematoxylin and eosin (H&E) or Prussian blue staining, respectively (Fig. 1f).

Immunostaining with a pan-Aβ polyclonal antibody S97 and anti-mouse IgG2a confirmed 3D6 localization to CAA-positive cerebellar meninges, hippocampal and cortical large vessels (arrows), and occasionally the choroid plexus, a conduit between blood and CSF (arrowhead, Supplementary Fig. 1a). Amylo-Glo, Collagen IV, and anti-Mouse IgG2a staining further demonstrated selective 3D6 binding along the vascular basement membrane (arrowheads) with very limited parenchymal labeling (Supplementary Fig. 1b). Quantitative analysis in the cerebellum confirmed preferential vascular over parenchymal plaque binding (paired t-test, *p* < 0.05; Fig. 1g,h). Bulk cerebellar qPCR revealed elevated *C4b* and *C3* mRNA levels in 3D6-treated mice compared with controls (Mann–Whitney, *p* < 0.05; Supplementary Fig. 1c), supporting early complement activation. Collectively, these results demonstrate that 3D6 initially binds to vascular amyloid, inducing early C1q deposition and complement activation.

### Short-term anti-Aβ 3D6 immunization promotes CAA-associated classical complement activation and vascular changes

We next examined whether short-term 3D6 immunization triggers classical complement activation and/or vascular damage by immunizing 16.5-mo-old *APP/PS1dE9;hApoE4* mice weekly with 500 µg 3D6 or IgG2a isotype control (i.p.) for 7 weeks (Fig. 2a). Previously, we showed that these mice develop an age-associated increase in CAA load [33]. Prussian blue staining revealed numerous microhemorrhages in the cortical and cerebellar meningeal and penetrating arterioles (asterisks) of 3D6-treated mice compared to controls (e.g., cortical microhemorrhages shown in Fig. 2b). Quantification confirmed a significant increase in hemosiderin deposits in 3D6-treated mice (Mann–Whitney, *p* = 0.0006; Fig. 2c). Amylo-Glo analysis showed a significant reduction in total cortical amyloid burden after 3D6 treatment (F1,12 = 35.43, p<0.0001, Two-way ANOVA) indicating a significant reduction in amyloid burden in the cortex (unpaired t-test, df = 14, t = 5.35, p = 0.0001; Fig. 2d) that was not sex-specific. Eosin staining revealed RBCs near cortical hemosiderin deposits only in 3D6-treated mice (Fig. 2e), consistent with RBC extravasation.

**Fig. 2:**
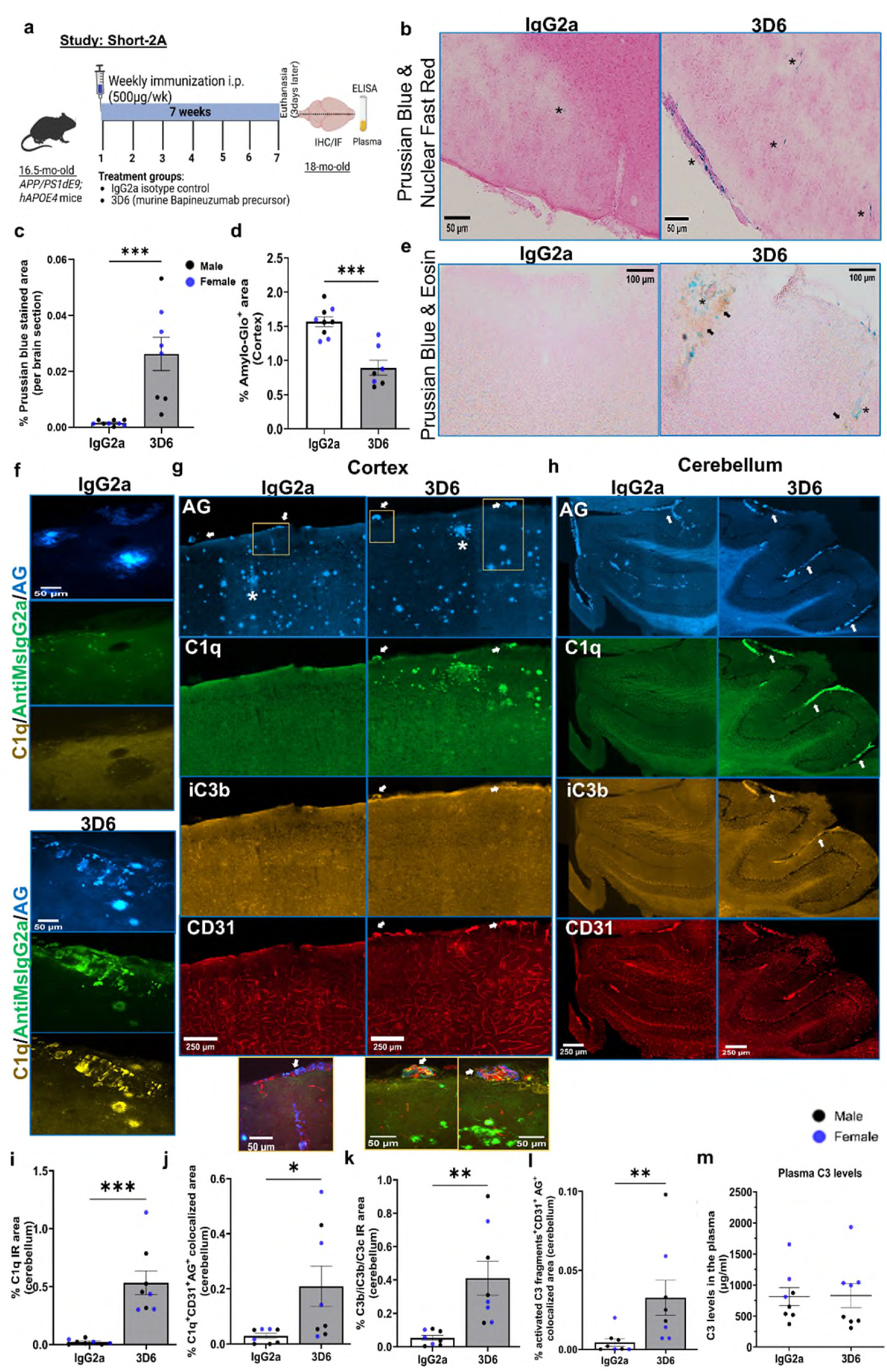
Short 7-week anti-Aβ 3D6 passive immunization reduced amyloid burden and induced deposition of CAA-associated complement C1q and activated C3 fragments and microhemorrhages without altering peripheral C3 levels. **a,** Schematic of 7-week treatment in 16.5-month-old *APP/PS1dE9;hAPOE4* mice (n = 7-8/group) with 3D6 or IgG2a control mAb. **b,** Prussian blue-labeled hemosiderin deposits in cortex indicating microhemorrhage (asterisks). Scale bar: 50 µm. **c,** Quantification of Prussian blue–positive area (%) in the whole sagittal brain section; Mann-Whitney test, p < 0.0005. **d,** Amylo-Glo (AG)–positive fibrillar amyloid area (%) in cortex; unpaired t-test, p < 0.0005. **e,** Prussian blue staining with eosin counterstain in cortex showing hemosiderin deposits (asterisks) and red-orange cytoplasm of extravasated RBCs (arrows). Scale bar: 100 µm. **f,** Immunostaining with anti-mouse IgG2a fluorescently-tagged secondary antibody, C1q and AG in brain sections from 3D6- and IgG2a-treated mice. Scale bar: 50 µm. Representative immunofluorescence staining in the **g,** cortex and **h,** cerebellum showing C1q and activated C3 fragments C3b/iC3b/C3c deposition near AG-positive plaques (asterisks) and vascular amyloid (arrow), Scale bar in **g & h**: 250 µm. Quantification of % immunoreactivity (IR) for **i,** C1q; **j,** C1q^+^CD31^+^AG^+^ colocalized area; **k,** activated C3 fragments C3b/iC3b/C3c; and **l,** activated C3 fragments^+^CD31^+^AG^+^ colocalization in cerebellum. **m,** Levels of complement C3 in the plasma following 7-weeks of passive immunization. Unpaired t-test in **j, k** and Mann-Whitney test for **i, l**. *p<0.05 and **p<0.005, ***p<0.0005. Data are expressed as mean ± SEM.

Using anti-mouse IgG2a as a surrogate for 3D6, we observed abundant antibody binding to both CAA and plaques, accompanied by C1q labeling (Fig. 2f). To assess complement activation more comprehensively, sections were co-labeled for Amylo-Glo, C1q, activated C3 fragments (C3b/iC3b/C3c), and the endothelial marker CD31. C1q deposition surrounded CAA-affected vessels and plaques in cortex and cerebellum (arrows, Fig. 2g,h). Quantitative analysis of the cerebellum, where 3D6-induced microhemorrhages showed significant increase in C1q (*p* = 0.0002; Fig. 2i) and C3b/iC3b/C3c immunoreactivity (*t* = 3.497, *p* = 0.0036; Fig. 2k) along pial vessels in 3D6-treated mice versus controls. Large CAA-positive vessels frequently displayed robust C1q and C3 labeling (arrows, insets Fig. 2g,h).

To quantify CAA-associated complement activation, colocalization between Amylo-Glo, CD31, and complement signals was determined. CAA-associated C1q (*t* = 2.429, *p* = 0.029; Fig. 2j) and C3b/iC3b/C3c (*t* = 2.479, *p* = 0.0265; Fig. 2l) deposition were significantly elevated in 3D6-treated mice. These results confirm complement activation along the cerebrovascular amyloid interface after short-term immunization. Plasma C3 levels were similar between treatment groups (Fig. 2m). Interestingly, plasma C3 levels were significantly higher in females than males in both groups (p<0.05). Together, these findings demonstrate that short-term 3D6 treatment promoted robust amyloid clearance but concurrently induced CAA-associated complement activation and microhemorrhage formation, supporting a link between cerebrovascular antibody binding, complement engagement, and early ARIA-like pathology.

### Effects of anti-Aβ mAb immunization on the brain and endothelial transcriptome

Following our observation of 3D6-induced microhemorrhages in the cerebellum of aged *APP/PS1dE9;hAPOE4* mice, we next examined transcriptomic changes in the same region to gain insight into mechanisms underlying ARIA. Twenty-month-old plaque- and CAA-rich *APP/PS1dE9;hAPOE4* mice were treated weekly for 7 weeks with 500 μg of anti-Aβ 3D6 or IgG2a isotype control mAb (n = 3 per group, Fig. 3a). PCA revealed clear separation between treatment groups (PC1 = 77%, PC2 = 11%, plot not shown). Differential expression analysis using DESeq2 identified 78 genes (raw p < 0.0005), of which 33 remained significant after multiple testing correction (padj < 0.05, Benjamini–Hochberg), including 30 upregulated and 3 downregulated genes in 3D6-treated mice (Fig. 3b,c). The upregulated genes included: *Mmp12, C3, Gpnmb, Gfap, Tmem176a, Gpr88, Ighm, Lyz2, Adcy7, C4b, Fcgr2b, Panx1, Adap2, AB124611, Igsf6, C1qa, Heyl, Csf3r, C1qb, Ctss, Nrros, Lgals3, C1qc, Pirb, Itgb2, Neu4, Ly9, Lag3, Arhgap45, Spag6,* and down-regulated genes included *Kcnq1ot1, Cecr2, Cyp2f2*. Although the sample size was low, the lone female mouse in this sub-study displayed a strong treatment effect, possibly due to higher amyloid burden commonly observed in female *APP/PS1dE9;hAPOE4* mice [33]. Gene set enrichment analysis showed upregulation of complement activation (*C1qa, C1qb, C1qc, C3, C4b*), glial activation (*Gfap*), IgG receptor signaling (*Fcgr1–3*), lysozyme activity (*Lyz2*), and lectin pathway genes (*Lgals3*) (Fig. 3d,e).

**Fig. 3:**
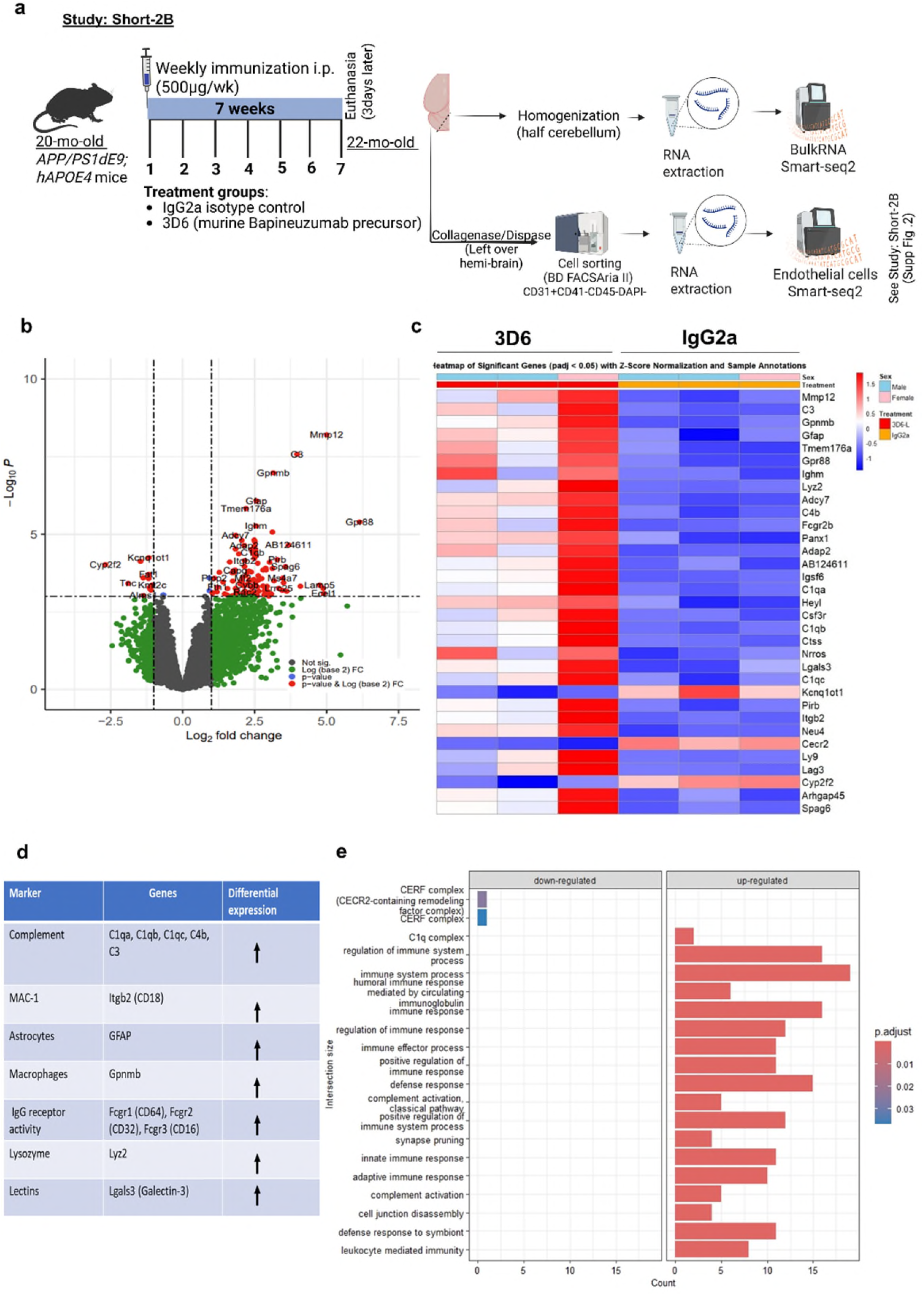
Passive anti-Aβ immunization for 7 weeks was associated with differential gene expression in the brain, particularly upregulation of inflammatory genes, including complement genes. **a,** Schematic for anti-Aβ immunotherapy in the 20-month-old *APP/PS1dE9;hApoE4* mice immunized for 7-weeks, 500 µg/week, followed by RNA extraction from cerebellar tissue and Smart-seq2 (n=3, 2M and 1F). **b,** Volcano plot showing differentially expressed genes with p-values < 0.0001. **c,** Pheatmap plot showing significantly expressed genes after multiple correction, p-adjusted value < 0.05. **d,** List of selected differentially expressed upregulated genes with respective functions. **e,** Bar plots with the top 20 up-regulated and down-regulated biological pathways.

To evaluate vascular-specific effects, endothelial cells (CD31⁺CD41⁻CD45⁻DAPI⁻) were FACS-sorted from the left hemibrain including cerebellum (Supplementary Fig. 2a). PCA again revealed separation between 3D6- and IgG2a-treated groups (PC1 = 42%, PC2 = 22%). Analysis identified 467 DEGs (p < 0.05), of which 23 remained significant after correction (padj < 0.05): The upregulated genes include *Lrg1, Abca1, Lcn2, Il4ra, Ptgs2, Tubb6, Itga5, Mir22hg, Vwf, Il1r1, Ch25h, Cd9, Runx1, Vmp1, Cfh, Kbtbd11, Cxcl2, Pim1, Tgif1,* and downregulated genes include *Cxcl12, Car4, Ndnf, Hmcn1* (see Supplementary data for full details). Enrichment analysis indicated downregulation of cell adhesion and extracellular matrix pathways and upregulation of stress, *NF-κB, IL-17,* and *PDGF* receptor signaling (Supplementary Fig. 2d). Notably, *Cfh*, which encodes complement factor H, a soluble glycoprotein that inhibits complement activation and promotes C3b degradation [34] was upregulated, suggesting a compensatory response to complement activation. qPCR from a small portion of cerebellar tissue validated the increased expression of *C1qc, C4b, C3, Lag3, Gfap, Gpnmb, Ch25h, Itgb2,* and *Lrg1* in 3D6-treated mice. Expression of *Tjp1* (*ZO-1*) was unchanged but *Ocln* was reduced, indicating subtle tight-junction changes, and *Cd59b*, a complement inhibitory gene, was reduced (Supplementary Fig. 3). Together, these findings demonstrate that anti-Aβ immunization induced robust transcriptomic changes involving complement activation, glial and endothelial stress, and immune signaling, processes that may collectively contribute to vascular fragility and ARIA-like pathology.

### MRI revealed microvascular lesions following chronic 3D6 immunization in mice

Anti-Abeta immunotherapies have been associated with ARIA, detectable by MRI. These side effects are more common in *APOE4* carriers [4]. In our initial MRI standardization protocol, we performed a 7-week passive immunization in 17-mo-old *APP/PS1;hApoE4* mice with 500μg/week of 3D6 mAb. We adopted 3D SWI_Fc FLASH and FLAIR sequences to identify any hypointense spots (microhemorrhages, ARIA-H) and fluid accumulation/edema (ARIA-E), respectively. Little if any signal was observed (data not shown). We increased dosing to 13 weeks of 500 μg/week 3D6 starting at 17 months of age (as shown in Fig. 4a) and detected noticeable hypointense spots (ARIA-H) in 20-mo-old *APP/PS1;hAPOE4* mouse brains seen by the T2*-FLASH-weighted sequence; however, ARIA-E remained undetectable. Hypointense spot areas indicative of microhemorrhages were confirmed on sagittal brain sections using Prussian blue hemosiderin staining (Supplementary Fig. 4). Nineteen slices covering the whole brain were analyzed in PBS controls and 3D6-treated mice. Cerebral microbleeds were significantly higher in 3D6-treated mice compared to age-matched PBS-treated controls (Unpaired t-test, df = 5, t = 7.295, p = 0.0008, Fig. 4b,c). Perls Prussian blue staining revealed minimal to no hemorrhages in PBS and IgG2a isotype control-treated mice, however, 3D6-treated mice had frequent hemosiderin deposits in the cortex, penetrating arterioles, hippocampal fissure, thalamus and cerebellum (arrowheads, Fig. 4e). Quantification revealed a significant increase in hemosiderin-positive staining in 3D6-treated compared to IgG2a isotype control-treated mice (Unpaired t-test, df = 8, t = 4.33, p = 0.0025, Fig. 4d). PBS- and IgG2a-treated mice showed similar results indicating that IgG2a isotype control treatment did not induce microhemorrhages.

**Fig. 4:**
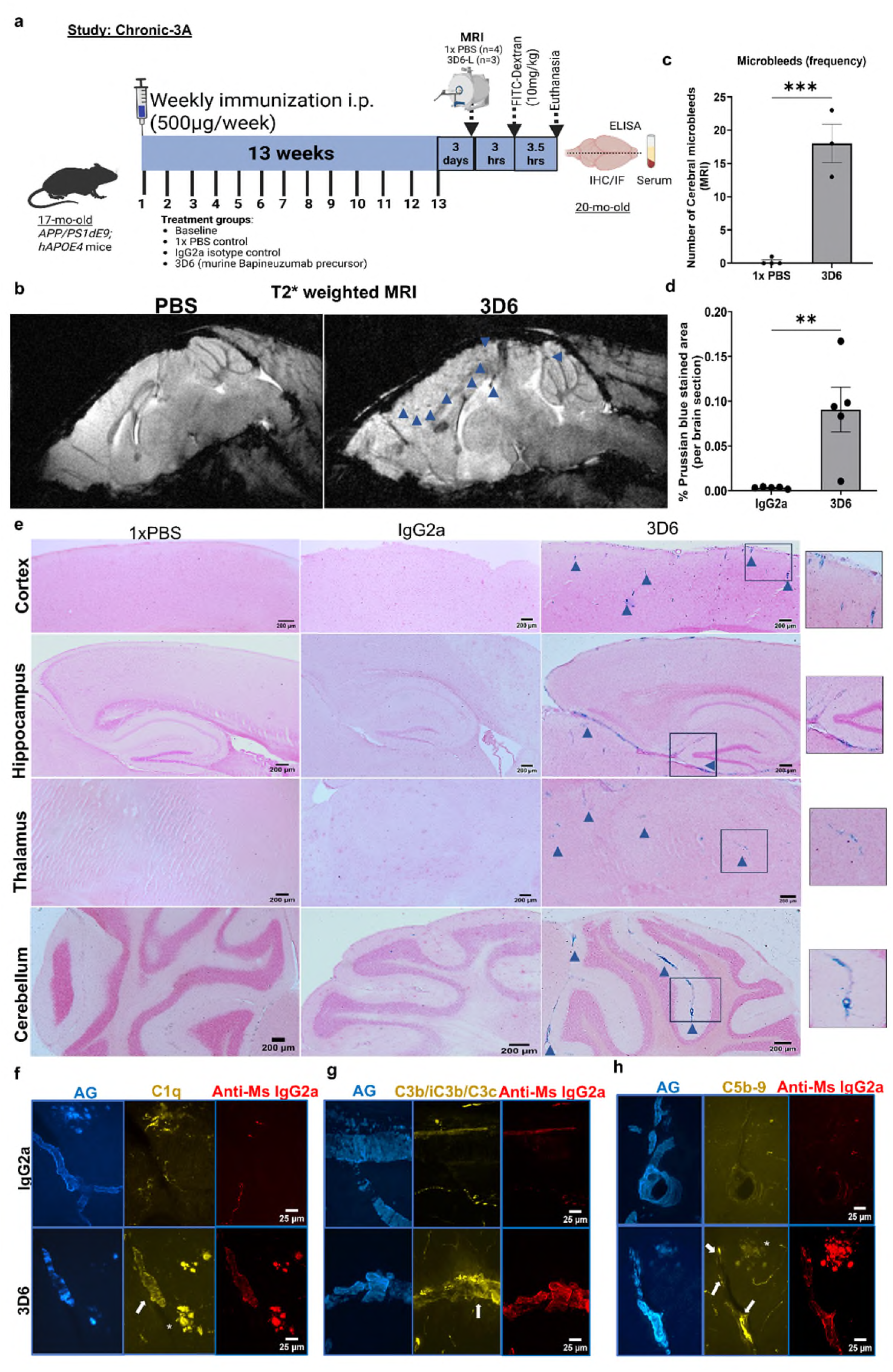
MRI and histopathology-based detection of immunotherapy-associated microbleeds and CAA-associated classical complement activation in mice chronically treated for 13 weeks. **a,** Schematic of anti-Aβ immunization with 500 µg/wk 3D6 IgG2a mAb and an IgG2a isotype control for 13 weeks from 17 to 20 months in aged male *APP/PS1dE9;hAPOE4* AD-like mouse model. For baseline data, baseline control mice were used. For spontaneous hemorrhage detection, 1x PBS-injected aged controls were used. **b,** T2*-weighted FLASH MRI of 20-month-old male *APP/PS1dE9;hApoE4* mice after 13 weeks of passive immunization with 3D6 mAb (500 µg/week), showing cerebral microbleeds as hypointense spots (arrowheads), compared to 1× PBS–treated aged controls. **c,** Frequency of cerebral microbleeds detected by T2* MRI in 3D6–treated (n = 3) and PBS-treated (n = 4) mice. **d,** Quantification of Prussian blue–positive (% hemosiderin-stained) area in IgG2a and 3D6–treated mice. Unpaired t-test: **p < 0.005, ***p < 0.0005. Data represent mean ± SEM. **e,** Representative Prussian blue–stained sections showing hemosiderin deposits (arrowheads, with selected examples in insets) in cortex, hippocampus, thalamus, and cerebellum from 1× PBS (n=4), IgG2a isotype control (n = 5), and 3D6 IgG2a–treated mice (n = 5). Scale bar: 200 µm. **f,** Immunofluorescence staining in the cerebellum showing 3D6 mAb localization (anti-Ms IgG2a) and C1q deposition at sites of vascular and parenchymal amyloid labeled with Amylo-Glo (AG). **g,** Detection of activated complement fragments C3b/iC3b/C3c at vascular amyloid sites associated with 3D6 mAb. **h,** Localization of membrane attack complex (MAC/C5b-9) along vascular amyloid (arrows) and plaques (asterisk) in 3D6–treated mice. Minimal complement staining is observed in IgG2a-treated controls. Scale bars: f–h, 25 µm.

### Chronic anti-Aβ 3D6 immunization resulted in CAA-associated complement activation, elevated cerebral C3 levels, and reduced serum C3

To evaluate complement involvement in 3D6-associated vascular changes, brain sections from 20-mo-old *APP/PS1;hAPOE4* mice (after 13-week treatment with 500 μg/week 3D6 or IgG2a) were immunostained for complement markers. 3D6 mAb robustly bound plaques and vascular amyloid estimated by anti-mouse IgG2a staining (Fig. 4f–h), accompanied by marked deposition of C1q (Fig. 4f), activated C3 fragments (C3b/iC3b/C3c; Fig. 4g), and the terminal complement complex C5b-9 (Fig. 4h). Double-labeling showed C1q enrichment around vascular amyloid and plaques (Fig. 5a), with significantly greater C1q immunoreactivity (unpaired t-test, df = 8, t = 3.185, p = 0.0129; Fig. 5b) and amyloid-associated C1q colocalization (Unpaired t-test, df = 8, t = 9.325, p < 0.0001; Fig. 5c) in 3D6-treated mice. C1q also colocalized with CD31+ vasculature (unpaired t-test, df = 8, t = 4.395, p = 0.0023; Fig. 5d).

**Fig. 5:**
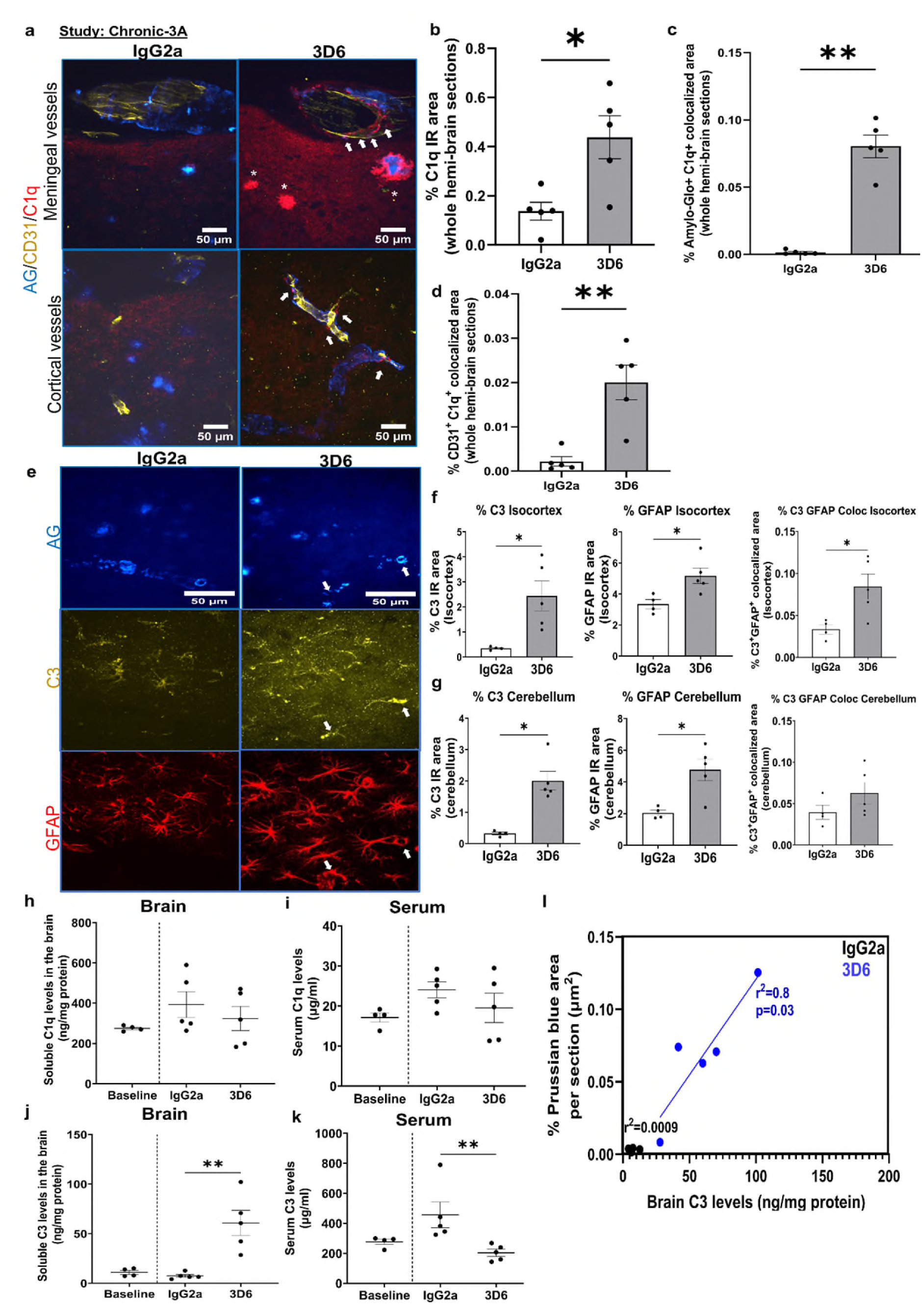
Chronic anti-Aβ passive immunization with 3D6 mAb was associated with elevated C3 levels in the brain and a corresponding lowering in the serum, in aged 20-mo-old male *APP/PS1dE9:hAPOE4* mice following 13 weeks of passive immunization. **a,** Immunostaining for fibrillar amyloid (Amylo-Glo, AG), endothelial marker CD31, and complement C1q in cortical and meningeal vessels. Asterisks indicate plaque-associated C1q immunoreactivity (IR); arrows indicate C1q accumulation along vessels. **b–d,** Quantification of (**b**), C1q IR (**c**), AG⁺C1q⁺ colocalized area and (**d**), CD31⁺C1q⁺ colocalized area. **e,** Representative images from the cortex showing complement C3 immunoreactivity in GFAP-positive astrocytes, including perivascular localization (arrows). **f–g,** Quantification of % area for C3, GFAP, and C3⁺GFAP⁺ colocalization in the (**f)** isocortex and (**g**) cerebellum. **h–i,** Soluble C1q levels in **h,** T-per brain homogenates and **i,** serum. **j–k,** Soluble C3 levels in **j,** T-per brain homogenates and **k,** serum measured by ELISA. **l,** Linear regression between brain soluble C3 levels and % Prussian blue–stained area (hemosiderin/microhemorrhages) across IgG2a and 3D6–treated groups. Unpaired t-test: *p < 0.05, **p < 0.005. Data represent mean ± SEM; n = 5 per group. Scale bars: a, e = 50 µm.

Cortical astrocytic C3 staining was increased after 3D6 treatment (Fig. 5e), with elevated C3 (unpaired t-test, df = 7, t=3.1, p=0.01), GFAP (unpaired t-test, df = 7, t=2.94, p=0.02), and C3+GFAP+ colocalization (unpaired t-test, df = 7, t= 2.87, p=0.02; Fig. 5f). We also observed increased immunoreactivity for C3 (Mann-Whitney test, p=0.01) and GFAP (unpaired t-test, df = 7, t= 3.456, p=0.01) and non-significant trend for increased C3+GFAP+ colocalization (p=0.2) in the cerebellum (Fig. 5g). CD206+ perivascular macrophages were modestly increased and showed strong C3 colocalization in cortex (unpaired t-test, df = 7, t= 9.976, p<0.0001) and cerebellum (df = 7, t= 6.087, p=0.0005; Supplementary Fig. 5c–f). While C1q ELISA levels in brain and serum remained unchanged (Fig. 5h,i), soluble C3 in brain homogenates rose ∼6-fold (F2,11 = 14.49, p = 0.0008, One-way ANOVA) and were significantly higher than baseline (Unpaired t-test, df =7, t= 3.449, p=0.0107) and isotype controls (Unpaired t-test, df =8, t = 4.198, p=0.0013; Fig. 5j). Conversely, serum C3 was reduced (F = 5.734, p = 0.0197; Mann–Whitney p = 0.0079; Fig. 5k), suggesting redistribution from periphery to brain or C3 cleavage to its fragments. Brain C3 levels correlated positively with Prussian-blue-defined microhemorrhages (R² = 0.80, p = 0.03; Fig. 5l).

3D6 immunization also enhanced C5b-9 activation (Fig. 6a). Quantification showed increased deposition of activated C3 fragments, C3b/iC3b/C3c (unpaired t-test, df =7, t = 9.07, p<0.0001), C5b-9 (unpaired t-test, df =7, t = 4.67, p=0.0023) (Fig. 6b,c). CAA vessel analysis (22–25 vessels/mouse) showed greater CAA+iC3b+ (unpaired t-test, df =8, t = 2.83, p=0.02) and CAA+C5b-9+ (unpaired t-test, df =6, t = 4.80, p=0.003) colocalization (Fig. 6d,e), while serum C5b-9 levels were elevated (t = 2.59, p = 0.04; Fig. 6f). CAA burden was modestly reduced (t = 1.94, p = 0.087; Fig. 6g). Histology revealed iC3b and Prussian blue overlap (arrows) with attenuated S97 Aβ signal (arrowheads, Fig. 6h), suggesting complement-mediated microvascular injury. Isotype controls showed strong vascular Aβ but minimal complement deposition or microhemorrhage.

**Fig. 6:**
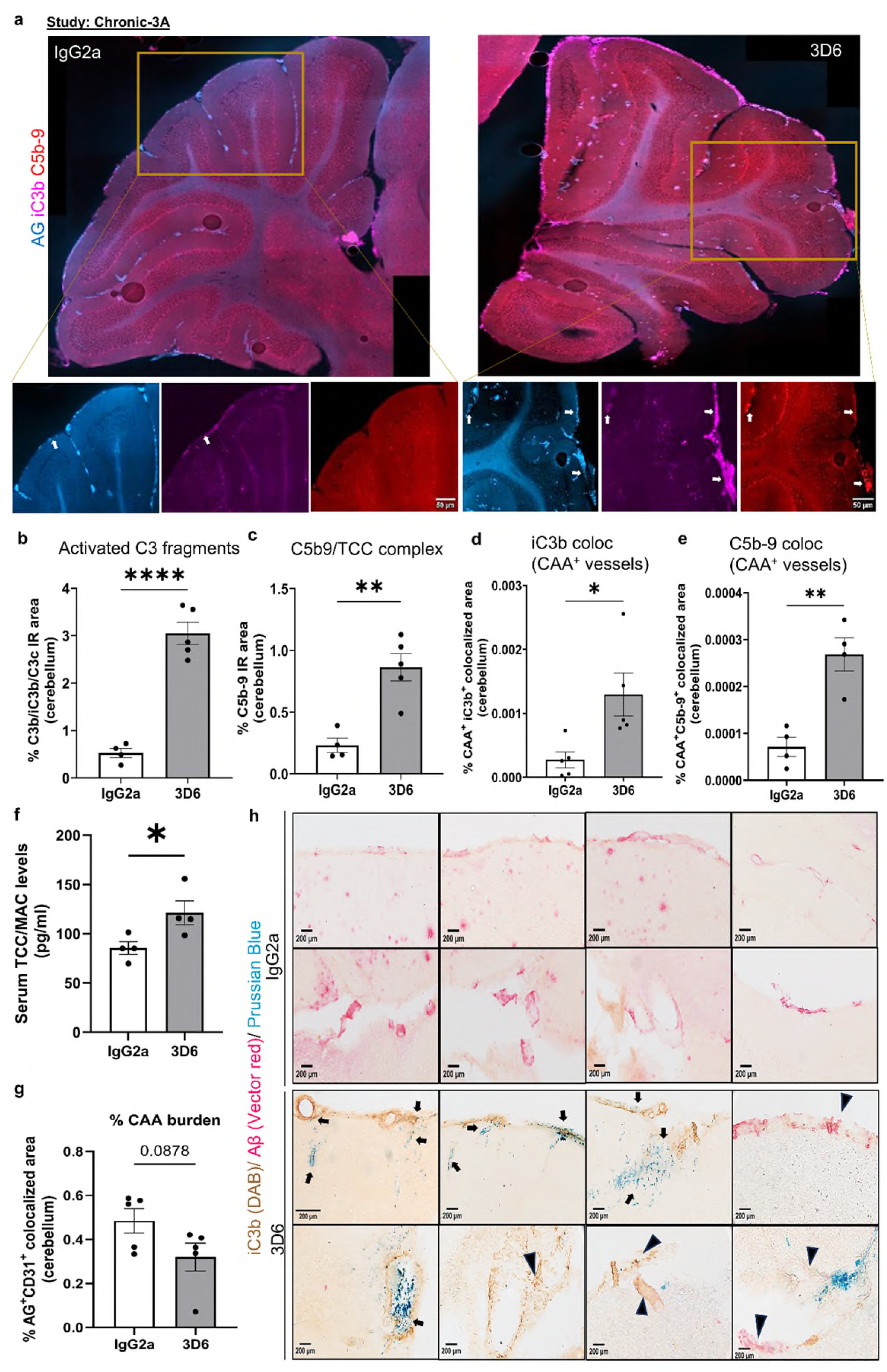
Chronic 3D6 mAb with 500 µg/wk was associated with increased activated C3 fragments and TCC/MAC complex in CAA-associated vessels. **a,** Representative immunofluorescence staining in the cerebellum area showing activated C3 fragments (C3b/iC3b/C3c) and TCC/MAC complex (C5b-9) along with fibrillar amyloid (Amylo-Glo dye, AG) in IgG2a isotype and 3D6 mAb passively immunized mice. Arrows indicate iC3b or C5b-9 immunoreactivity along CAA^+^ vessels stained with AG. Scale bar 50 µm. Quantification of % area immunoreactivity for **b,** iC3b **c,** C5b-9 per brain section. Approximately 22 to 25 Amylo-Glo^+^ CAA vessels were outlined to quantify **d,** CAA^+^iC3b^+^ colocalization & **e,** CAA^+^C5b9^+^ colocalization area. **f,** Serum TCC/MAC levels measured by ELISA **g,** % CAA estimated by AG^+^CD31^+^ colocalization data. **h,** Representative double IHC staining for iC3b (DAB), S97 Aβ (Vector red), with Prussian blue combination for microhemorrhages detection in the 3D6 treated mice showed iC3b-associated microhemorrhages in leptomeningeal or penetrating vessels (arrows) and signs of vascular amyloid clearance (arrowheads) in vessels with complement deposition. In IgG2a-treated mice, vascular amyloid/plaque staining is prominent with minimal complement and Prussian blue staining (arrows), Scale bar=20 µm. N=5/group, Parametric unpaired t-test, *p<0.05 and **p<0.005, ****p<0.00005. Data are expressed as mean ± SEM.

### Chronic anti-Aβ 3D6 immunization increased BBB leakiness and MMP expression in CAA-laden vessels

To assess BBB integrity after chronic 3D6 treatment, the same 17-mo-old *APP/PS1;hAPOE4* (treated with 500 μg/week 3D6 or IgG2a) were administered a lysine fixable FITC-Dextran injection 3.5 h before euthanasia (Fig. 4a). Anti-mouse IgG2a staining confirmed mAb binding to plaques and vascular amyloid (unpaired t-test, df=8, t=7.6, p<0.0001, Fig 7a, bar chart not shown). 3D6-treated mice displayed pronounced MMP-9 labeling along vascular amyloid (Fig. 7a) that reached significance (t = 7.458, p < 0.0001; Fig. 7c). FITC-Dextran leakage around CAA+ vessels was increased (t = 4.66, p = 0.0016; Fig. 7d) and strongly colocalized with MMP-9 (t = 4.12, p = 0.003; Fig. 7e) confirming BBB compromise. Enhanced vWF was seen around CAA vessels (Suppl Fig. 5a), suggesting endothelial stress or vascular injury. Leakage in IgG2a controls likely reflects age and CAA-related APOE ε4-linked BBB dysfunction and impaired perivascular clearance [35,36]. Thus, chronic 3D6 immunization exacerbated CAA-associated MMP-9 accumulation, FITC-Dextran leakage, and vWF deposition, consistent with vascular damage. This is consistent with our 7-week 3D6 treatment finding of upregulation of Matrix metalloproteinase MMP-12 gene expression, which is known to correlate with cortical hemorrhage [37,38].

**Fig. 7:**
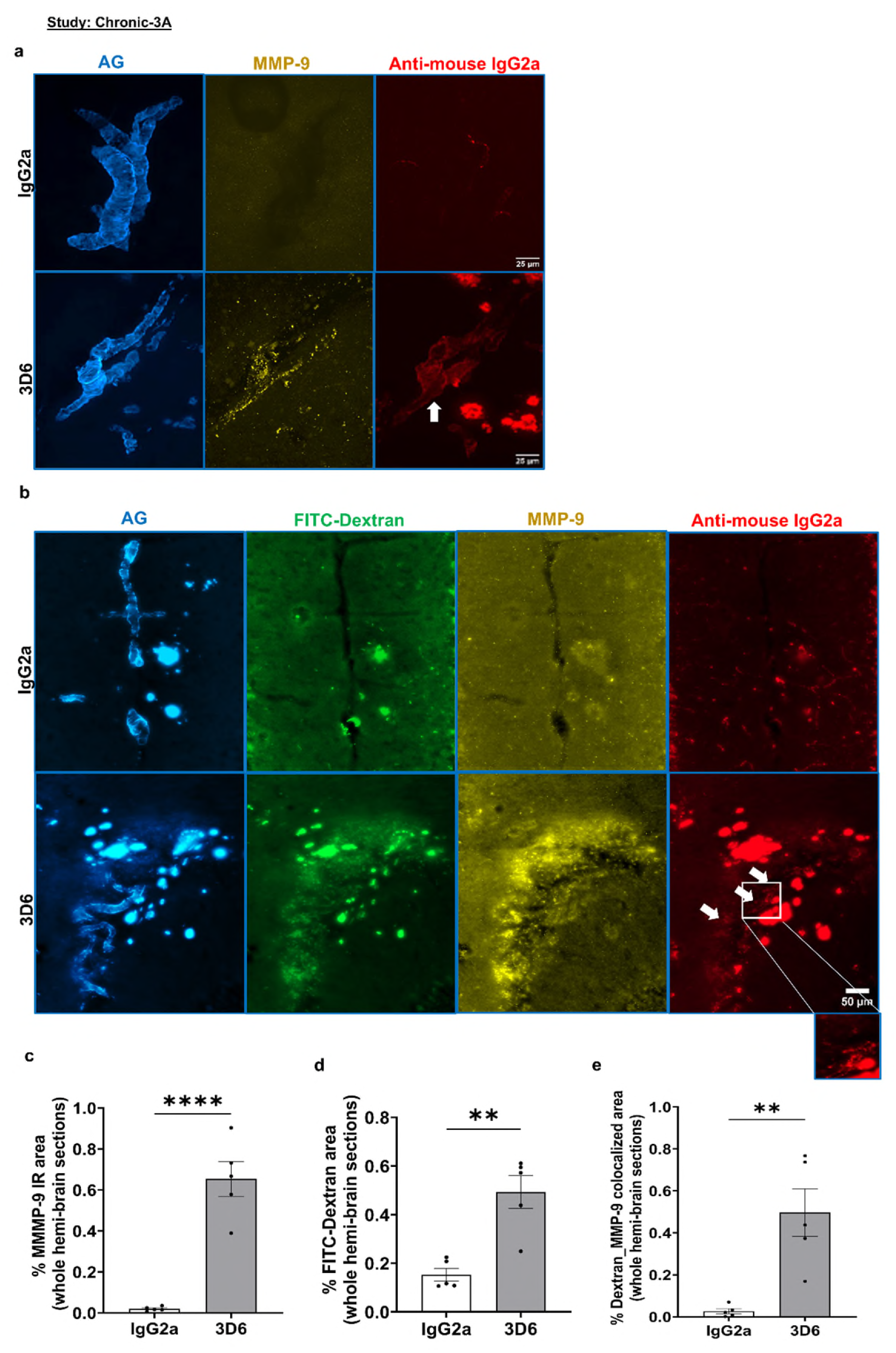
Chronic anti-Aβ passive immunization with 500 µg/wk of 3D6 mAb was associated with mAb-associated matrix metalloproteinase expression and FITC-Dextran leakage along the CAA-laden vessels. Following passive immunization, mice were injected (i.p.) with Dextran fluorescein (10 mg/kg body weight) and euthanized 3.5 hours later. **a,** Representative 40× immunofluorescence images showing probable 3D6 mAb binding (anti-Ms IgG2a) to vascular amyloid (arrows) and plaques (Amylo-Glo, AG) in the cerebellum, with matrix metalloproteinase-9 (MMP-9) expression. **b,** Slide-scanned 20× images showing AG, FITC-Dextran, MMP-9, and anti-Ms IgG2a staining. Arrows and inset highlight CAA-associated antibody colocalization. **c–e**, Quantification of **c,** MMP-9 immunoreactivity per brain area, **d,** FITC-Dextran deposition, and **e,** FITC-Dextran and MMP-9 colocalized area. Unpaired t-test: **p < 0.005, ***p < 0.0005, ****p < 0.00005. Data represent mean ± SEM; n = 5 per group. Scale bar: a=25 µm, b=50 µm.

### Chronic anti-Aβ passive immunization with 3D6 mAb reduced cerebral amyloid levels in *APP/PS1;hAPOE4* mice

To examine amyloid clearance, cerebral Aβ levels were measured from the same mice treated for 13 weeks with 500 μg/week 3D6 or IgG2a (Fig. 4a). Guanidine-soluble Aβ42 differed by treatment (F2,11 = 6.937, p = 0.0112, One-way ANOVA) and was significantly lower in 3D6- vs. IgG2a isotype control-treated mice (unpaired t-test, p < 0.005; Fig. 8a) with a nonsignificant ∼20 % decrease from baseline. Aβ40 was decreased (F2,11 = 18.80, p = 0.0003, One-way ANOVA) with 3D6-treated mice having significantly lower Aβ_40_ levels (unpaired t-test, p < 0.0005) while levels in IgG2a controls were higher than baseline consistent with age-related amyloid deposition (p < 0.005; Fig. 8b). The Aβ42/40 ratio rose after 3D6 treatment (p < 0.05; Fig. 8c) towards baseline levels suggesting preferential Aβ40 clearance.

**Fig. 8:**
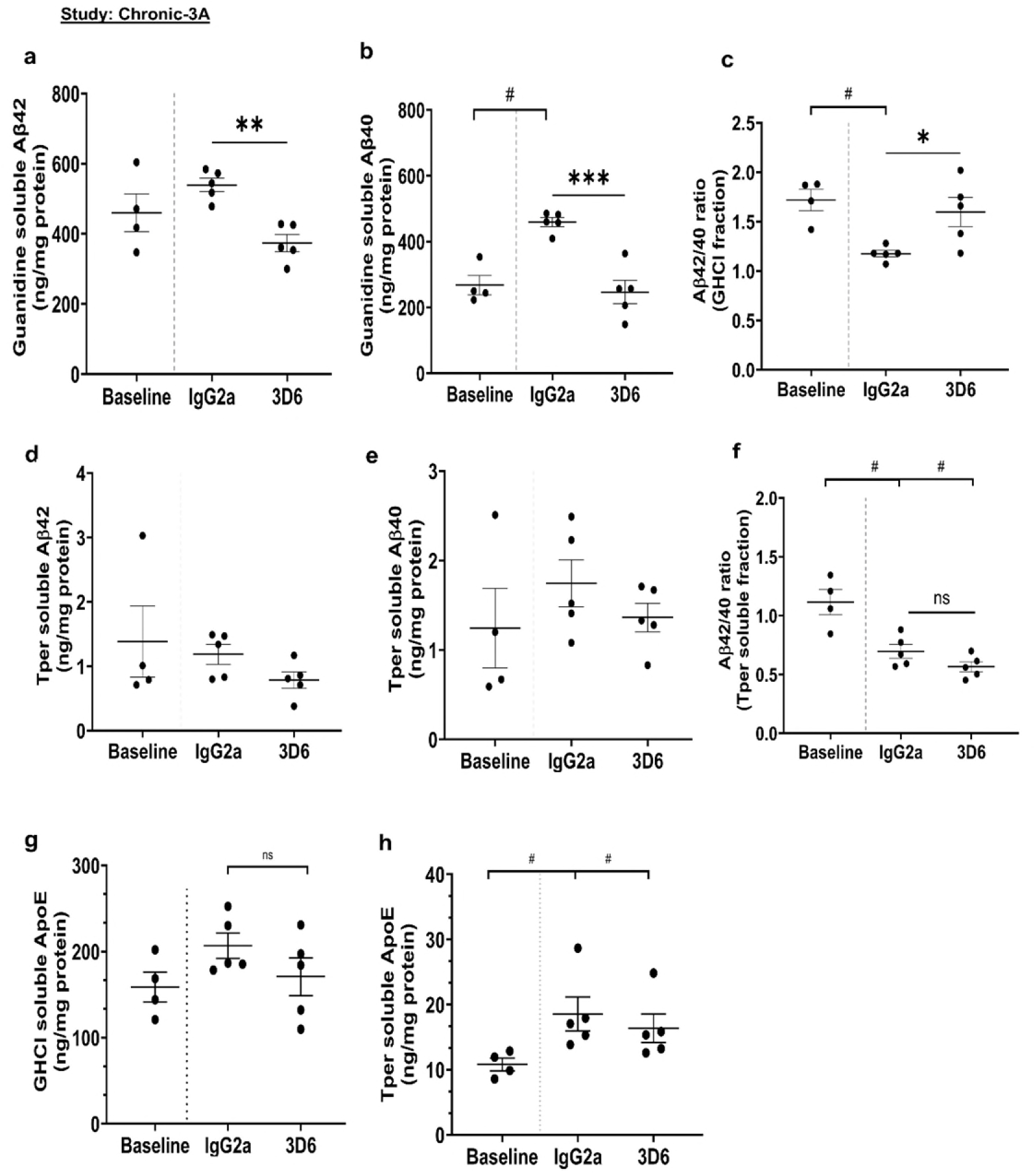
Chronic anti-Aβ passive immunization with 500 µg/wk 3D6, lowers amyloid levels in the brain. 3D6 IgG2a mAb and an IgG2a isotype control were administered for 13 weeks from 17 to 20 months in aged male APP/PS1dE9; hAPOE4 mice. For baseline data, baseline control mice were used. ELISA data for **a,** Aβ42 **b,** Aβ40 **c,** Aβ42/40 ratio in the Guanidine HCl brain fractions, and **d,** Aβ42 **e,** Aβ40 **f,** Aβ42/40 ratio in the Tper soluble brain extracts. Levels of apoE in the brain **g,** GHCl soluble extracts, and **h,** Tper soluble extracts. Data are expressed as mean ± SEM, n=4-5/group. Unpaired t-tests (IgG2a vs 3D6), *p<0.05, **p<0.005, ***p<0.0005 and #p<0.005 (for exploratory comparison between baseline vs IgG2a or 3D6 group). Data are expressed as mean ± SEM.

No significant differences were seen in T-per soluble Aβ42 and Aβ40 levels between groups (Fig. 8d,e). Both IgG2a- (p < 0.005) and 3D6- (p = 0.0005) treated mice had reduced T-per soluble Aβ42/40 ratios compared to baseline levels, consistent with increasing plaque deposition with age (Fig. 8f). ApoE levels in GHCl (Fig. 8g) and T-per fractions (Fig. 8h) remained largely unchanged, aside from an increase in soluble ApoE levels in treated groups vs baseline (p < 0.05). Overall, chronic 3D6 treatment decreased total Aβ burden without altering ApoE levels.

### Chronic anti-Aβ 3D6 immunization with a lower dose triggered CAA-associated complement and immune cell activation, and increased central but not peripheral C3 levels

To determine dose-dependence, 16-mo-old *APP/PS1;hAPOE4* mice received 350 μg/week 3D6 or IgG2a for 15 weeks (Fig. 9a). As expected, anti-mouse IgG2a colocalized with vascular amyloid and C1q deposition (Fig. 9b) which was associated with CD68+ Iba1+ microglia/macrophages (Fig. 9c). C3 levels were elevated in T-per brain homogenates in 3D6-treated mice (F1,13 = 12.66, p=0.0035, Two-way ANOVA) that was not specific for sex. Significantly higher cerebral C3 levels were seen in 3D6-treated mice (t = 3.93, p = 0.0013; Fig. 9d), consistent with our previous 7-week RNA-seq and 13-week brain C3 ELISA results. Plasma C3 was not significantly affected by 3D6 treatment (Fig. 9e). Thus, even a lower-dose 3D6 elicited CAA-linked C1q and C3 activation with immune cell recruitment, increasing brain but not peripheral C3 levels.

**Fig. 9:**
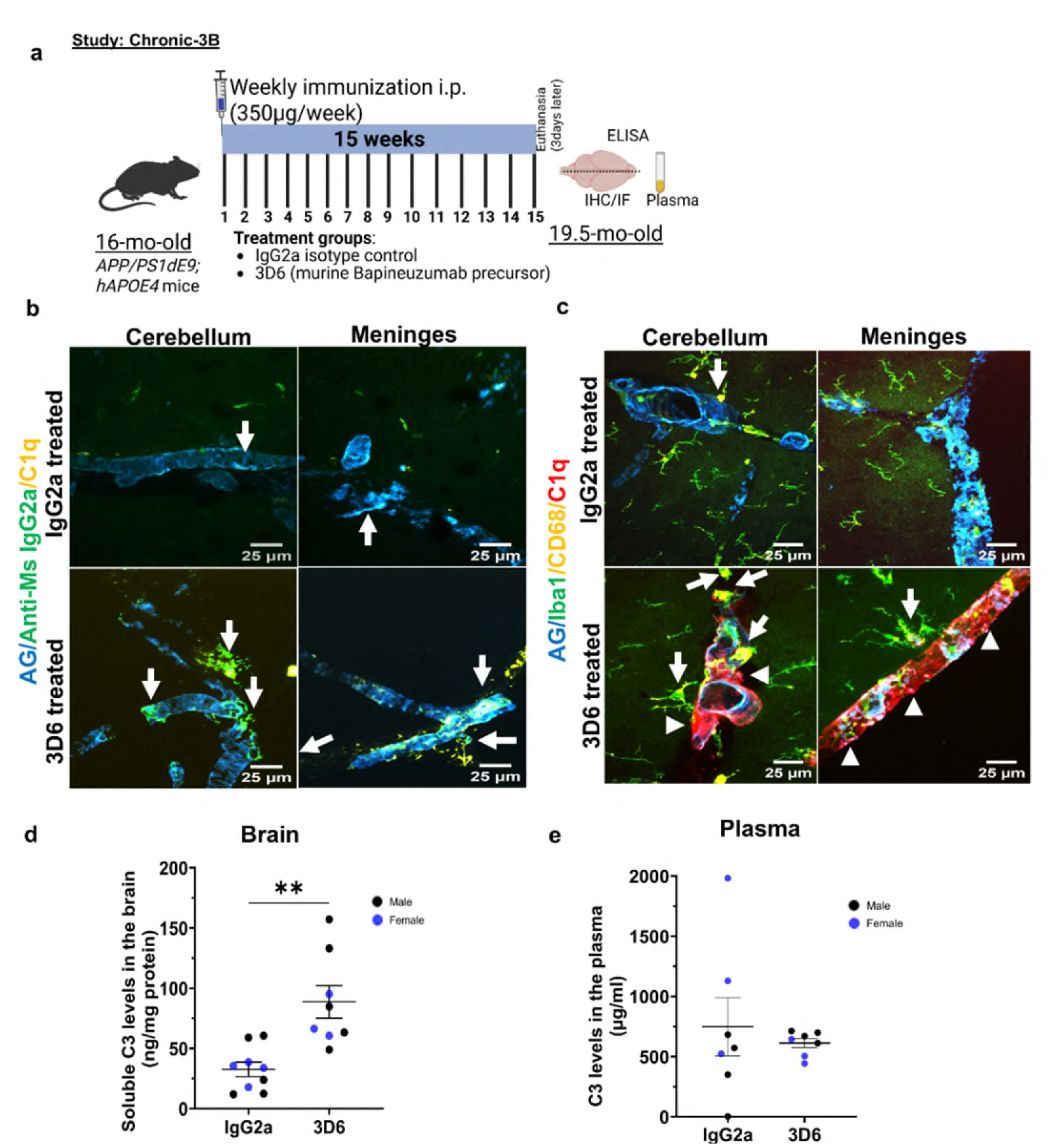
Chronic anti Aβ-immunization with 350 µg/wk of 3D6 mAb for 15 weeks was associated with mAb binding to vascular amyloid, complement, and immune cell activation and increased brain C3 levels but had no effect on blood C3 levels. **a,** Schematic of 15-week passive immunization in 16-month-old *APP/PS1dE9; hAPOE4* mice (n = 8–9/group) with 3D6 or IgG2a isotype control (350 µg/week). **b,** Immunostaining in the cerebellum and meningeal layers showing anti-Ms IgG2a (to detect mAb) binding, vascular amyloid (Amylo-Glo), and C1q. Arrows indicate co-localization of mAb, vascular amyloid, and C1q. **c,** Multiple immunolabelling with Iba1, CD68 showing amoeboid-like activated microglia/macrophages (arrows) near CAA-laden vessels (Amylo-Glo) and C1q staining in the cerebellum and meninges. Arrowheads indicate C1q accumulation in cerebellar and meningeal vessels. **d–e,** Soluble C3 levels measured in **d,** Tper brain extracts and **e,** plasma. Mann-Whitney test, **p < 0.005. Data are expressed as mean ± SEM. Scale bars: b–c, 25 µm.

## Discussion

ARIA, characterized by brain edema and effusion, superficial siderosis and microhemorrhages, has occurred in a subset of AD patients treated with certain anti-amyloid monoclonal antibodies (e.g., bapineuzumab, gantenerumab, aducanumab, lecanemab and donanemab), particularly in individuals carrying the APOE ε4 allele [25]. Clinical studies in humans have shown that ARIA-E typically appears early after the start of immunization with ARIA-H often occurring later as seen with aducanumab [10], lecanemab and donanemab [8,9]. Here, we investigated the timing and mechanisms underlying ARIA by examining the effects of acute, short-term and long-term passive immunization with the murine bapineuzumab precursor 3D6. ARIA risk is influenced by age, dose, APOE genotype, and pre-existing CAA [39–42], therefore we conducted our studies in aged *APP^NL-G-F^* KI (knock-in) and *APP/PS1dE9;hAPOE4* mice which develop CAA mostly in the cerebellar arteriolar leptomeninges and penetrating arterioles and to some extent in cortical arteriolar leptomeninges and penetrating arterioles. Upon immunization, 3D6 first bound CAA followed by plaque binding. With further dosing, mice developed ARIA-like features including RBC extravasation, BBB leakage, microhemorrhages, and associated immune and cellular changes.

In our acute single or two-dose paradigms in aged APP*^NL-G-F^* mice, 3D6 preferentially localized to vascular amyloid within large meningeal and penetrating vessels rather than parenchymal plaques. High-resolution imaging revealed that 3D6 accumulated along the basement-membrane side of CAA deposits in a striated pattern adjacent to the smooth-muscle layer, occasionally extending toward the endothelial lumen. In some regions, antibody labeling spread from CAA-laden vessels to adjacent surrounding areas and, infrequently, plaques, suggesting perivascular transport of the antibody through interstitial drainage pathways [43,44]. These findings imply that circulating anti-amyloid mAbs can access vascular amyloid either directly from the bloodstream via the lumen or via perivascular routes soon after administration. These vascular binding patterns of the mAbs coincided with C1q deposition and activation of the classical complement cascade, accompanied by increased cerebellar *C4b* and *C3* mRNA expression in 3D6-treated mice after two doses, indicating early complement activation at the sites of vascular amyloid.

Given the known barrier compromise in CAA vessels [44], such complement-driven inflammation may precede and promote BBB leakage or vessel rupture [45], facilitating increased antibody entry, binding to CAA and vascular injury. Indeed, human IgG1 antibodies showing high affinity for CAA fibrils such as aducanumab, bapineuzumab, and gantenerumab are also those most frequently associated with ARIA-E, whereas, solanezumab which recognizes soluble Aβ, and crenezumab, which is on an IgG4 backbone, display weaker CAA binding [46] and lower ARIA rates [47]. Lecanemab binds to smaller diffusible aggregates or protofibrils with greater avidity compared to other Aβ species [48], which may be less prevalent in CAA, possibly explaining in part the lower ARIA rates in Leqembi-treated patients. Donanemab, which targets the N-terminal truncated and modified pGlu3-Aβ species, has higher ARIA rates [7], potentially due to the abundant presence of such truncated Aβ species in the vasculature [49]. Thus, differences in antibody-specific binding to vascular amyloid, especially early in treatment, may contribute to immunotherapy-associated vascular side effects, alongside active clearance by microglia and the triggering of an immune response.

Although no microhemorrhages or RBC extravasation were detected in our acute studies, robust C1q deposition along CAA-rich vessels indicated early initiation of complement signaling. Collectively, our data suggest that vascular amyloid constitutes a primary site of antibody engagement during initial infusions, triggering local complement activation that may predispose to ARIA during subsequent treatment cycles. This mechanistic link is highly relevant to human AD, where CAA occurs in ∼80% of cases [50] and, depending on severity, vascular amyloid may become exposed to the bloodstream [49]. Antibody binding to exposed amyloid in the blood vessel wall could initiate complement-mediated vascular inflammation and BBB damage, akin to CAA-related inflammation (CAA-ri). Proteomic analyses of human CAA microvasculature have indeed shown enrichment of secreted and membrane-bound complement proteins including C1q and C3 [51]. Overall, our data demonstrates that early binding of anti-Aβ antibodies to CAA triggers classical complement activation and therefore approaches bypassing this, for example, by exploiting transferrin-receptor–mediated transport to deliver antibodies through capillaries rather than amyloid-laden arteries might therefore mitigate ARIA risk [52] as suggested by the promising clinical trial results with Trontinemab.

Because of the strong clinical association between APOE ε4 and ARIA incidence [25,39–42], and APOE ε4’s known ability to form complexes with C1q [53] and CFH [54] that regulate complement activation, we next evaluated 3D6 immunization in aged *APP/PS1dE9;hAPOE4* mice. Short-term (7-week) weekly dosing at 500 µg reduced fibrillar amyloid but markedly increased vascular C1q, C3b/iC3b/C3c deposition, and complement-related gene expression. Transcriptomic analyses revealed upregulation of inflammatory and complement genes (*C3, C1q, C4b*), along with markers of astrocytosis (*Gfap*), macrophage activation (*Gpnmb*), lysosomal and lectin genes (*Lyz2, Lgals3*), and endothelial stress/vascular injury markers (*Lrg1, Ch25h, Lcn2*) [55–57]. Concomitant induction of *Abca1* and *Cfh*—genes involved in cholesterol efflux and complement regulation suggested compensatory mechanisms limiting inflammation [58,59]. This increase in complement signaling in our mouse model is corroborated by a recent study, which demonstrated dysregulated complement signaling in myeloid cells (macrophages and microglia) and upregulation of pathways related to vascular integrity and angiogenesis in lecanemab-treated patients [60]. Overall, the transcriptional changes observed likely reflect vascular stress and damage control responses, coinciding with RBC extravasation and hemosiderin accumulation detected histologically, indicative of microhemorrhages. Similar to our findings of increased CAA-associated C5b-9 deposition in long-term treated mice, a study in humans showed increased deposition of C5b-C9 in the vascular wall in leptomeningeal and penetrating CAA vessels in patients treated with Aducanumab [61]. Such vascular compromise may also increase BBB permeability and fluid accumulation [62], mimicking ARIA-E observed early in treatment with clinical antibodies [8,19,20], where edema and microhemorrhages can occur independently or concurrently in patients [25,63–65], underscoring the heterogeneous vascular response to Aβ immunotherapy. Therefore, in-depth proteomic profiling of blood may help identify endothelial biomarkers associated with vascular stress, enabling detection of patients at risk for ARIA and provide strategies to mitigate treatment-related vascular injury.

Complement activation downstream of C3 may further amplify vascular injury as evident in focal cerebral ischemia, cerebral injury and CNS vasculitis [66–68]. C3 cleavage generates C3b/iC3b for opsonization and clearance of immune complexes, and C3a, a potent anaphylatoxin promoting chemotaxis and inflammation via binding to C3aR [69]. In our studies, elevated vascular iC3b deposition near CAA associated with mAb-C1q activation likely contributed to vessel fragility and leakage [70–72]. This may drive vascular smooth muscle cell phenotypic switching or fibrous cap formation. Also, C3b engagement with CR3 on microglia can mediate amyloid phagocytosis but may also contribute to polarization of macrophages [73] around the vascular wall, causing a pro-inflammatory M1-like phenotype. Further, complement-immune opsonized complexes can trigger inflammasome activation, leading to a chronic inflammatory state [74]. Prior work has shown that endothelial C3a/C3aR signaling enhances VCAM1 expression, vascular inflammation, immune cell recruitment, and BBB dysfunction during aging or cerebral injury [67], whereas endothelial C3aR deletion confers protection [75]. Although C3aR is mainly expressed on microglia and neurons [69], ischemic or edematous conditions can induce its expression on endothelial cells [76,77]. Accordingly, we observed increased C3aR and C5aR expression in brain bulk RNA-seq data. Although endothelial C3aR upregulation was not clearly detected in our dataset and may need in-depth sequencing, increased C3aR/C5aR expression at the tissue level suggests dysregulated C3a-C3aR and C5a-C5aR axes, which may upregulate inflammation around CAA vessels and promote endothelial dysfunction which may, in turn, increase the risk of ARIA.

To extend our observations from acute and short-term paradigms showing early C1q and C3 deposition, we next examined whether prolonged 3D6 immunotherapy promotes vascular amyloid clearance and full complement activation, including formation of the terminal membrane attack complex (MAC; C5b-9). Using T2*-weighted MRI, we found that 13-week treatment with 500 µg 3D6 mAb induced cerebral microhemorrhages across multiple regions including the leptomeninges, cortex, hippocampal sulcus, thalamus, and cerebellum consistent with reports from antibody trials [4]. Edema (ARIA-E) was not detected using our FLAIR-MRI protocol, possibly reflecting insufficient resolution and the transient nature of ARIA-E compared with the more persistent ARIA-H [62]. Comparisons with age-matched PBS controls revealed negligible or no spontaneous microbleeds, and similar Prussian blue staining between PBS and IgG2a groups confirmed that the observed lesions were treatment-specific rather than age-related.

Long-term 3D6 dosing significantly reduced guanidine-soluble Aβ42 and Aβ40, with the Aβ42/40 ratio approaching baseline, indicating a shift toward earlier-stage pathology. Although changes in Tper-soluble Aβ were not significant, a downward trend in Aβ42/40 suggested differential effects on soluble versus insoluble pools. Given that CAA contains both Aβ42 and Aβ40 with Aβ40 as the dominant isoform [78] these reductions likely reflect enhanced clearance of vascular amyloid. Histological analyses supported this interpretation, showing attenuated or patchy CAA staining and colocalization of iC3b with eroded vascular amyloid, consistent with complement-mediated opsonization and clearance [79]. These features paralleled findings from other passive immunization studies [80], although some reports described increased vascular amyloid following treatment [81], likely due to differences in mouse models, antibodies and treatment paradigms. Notably, antibody-driven neutralization of parenchymal plaques can mobilize soluble Aβ toward the vasculature, where distinct kinetic profiles of perivascular drainage and complement activation [3,82] may govern the extent of vascular stress and microhemorrhage formation. Together, these results indicated that chronic 3D6 immunotherapy enhances complement-dependent vascular Aβ clearance while concurrently predisposing fragile CAA-laden vessels to microhemorrhages at sites of active opsonization.

Previously, CAA pathology has been associated with activation of complement factors C1q, C4, and C3, along with deposition of the terminal membrane complex C5b-9 in vessels prone to hemorrhage, highlighting the detrimental role of complement in vascular injury [28]. We hypothesized that chronic 3D6 immunization would exacerbate this cascade. Indeed, in long-term 3D6-treated mice, we observed increased deposition of C1q, C3 fragments (C3b/iC3b/C3c), and C5b-9 around mAb-bound plaques and CAA, particularly within the cerebellum. These complement components colocalized with anti-mouse IgG2a, a surrogate marker of 3D6, and amyloid deposits, supporting antibody-driven immune-mediated amyloid clearance [83]. Enhanced C3 and GFAP staining, and their colocalization within astrocytes, further indicated that reactive astrocytes are a major source of local complement C3 production [84,85]. C3 released from CAA-associated astrocytes may be cleaved to C3b which may bind CR3 receptors on recruited microglia, macrophages, and monocytes to promote phagocytosis and vascular amyloid clearance [86] or C3a which may engage endothelial C3aR to induce vascular inflammation, barrier compromise, and BBB dysfunction [75].

While microglia likely contribute to vascular inflammation, recent studies highlight a predominant role for border-associated macrophages (BAMs) in driving oxidative stress [86] and for perivascular macrophages (PVMs) and peripheral monocytes in promoting microhemorrhages [87,88]. Consistent with this, we observed increased CD206+ PVM/BAM populations and elevated C3+CD206+ colocalization in 3D6-treated mice, indicating active recruitment of macrophage-lineage cells to complement-activated sites. Notably, macrophage-derived migrasomes containing Aβ40 have been shown to carry pro-inflammatory molecules that exacerbate complement activation and BBB injury in CAA [89], an aspect not yet explored in passive immunotherapy-associated ARIA. Given that all complement pathways converge on C3, the positive correlation between brain C3 levels and hemosiderin deposition suggests that complement activation-driven inflammation is a key mediator of vascular injury. This interpretation is further supported by the elevated iC3b and C5b-9 immunoreactivity, their colocalization with CAA, and increased serum C5b-9 levels in 3D6-treated mice. Notably, although peripheral C3 levels decreased in these mice, an increase in circulating C5b-9 indicates robust terminal pathway activation, consistent with the other reports, showing increased circulating C5b-9 levels in both mouse models and human patients with CAA [89]. Similar to our findings, a recent study showed increased deposition of C5b-C9 in the vascular wall of leptomeningeal and penetrating CAA vessels in patients treated with Aducanumab [61]. Although peripheral C3 levels have been reported to be significantly elevated in CAA patients and may serve as a biomarker for CAA [90], short-term immunization with 3D6 in mice increased brain C3b deposition without altering plasma C3 levels, whereas long-term immunization led to increased brain C3 accompanied by reduced peripheral C3 levels, suggesting potential redistribution of peripheral C3 into the brain. This effect was dose-dependent and may reflect increased central complement demand and or BBB dysfunction during long-term immunization, highlighting C3 and C5b-9 as complementary biomarkers for diagnosing complement associated vascular injury and ARIA risk.

As indicated earlier, 3D6 immunization caused increased RBC extravasation following short-term immunization. Impairment in vascular integrity and BBB leakage were further validated by increased extravasation of peripherally-injected FITC-Dextran in antibody-bound amyloid-laden vessels, and by increased deposition of matrix metalloproteinases like MMP-9 in agreement with other anti-Aβ studies [87]. This is consistent with findings in human CAA patients where cerebrovascular MMP-9 imbalance is associated with intracerebral hemorrhages [91]. The colocalization of MMP-9 with vascular amyloid and anti-Aβ mAb further suggests that amyloid clearance at the vasculature may lead to extracellular matrix remodeling, BBB breakdown or vascular damage as shown by increased CAA-associated deposition of vWF in our 13-weekly immunized mice or increased endothelial *vWF* mRNA expression in our 7-weekly immunized mice, implicating a role of vascular inflammatory mediators and complement activation in modulating vascular permeability and edema formation [92]. Interestingly, FITC-Dextran leakage was also observed to some extent in age-matched IgG2a control mice, which could be due potentially to the influence of *APOE*ε4 on BBB integrity [35].

Overall, we have demonstrated that early 3D6 infusions preferentially targeted cerebrovascular amyloid rather than parenchymal plaques, and induced C1q deposition, initiating classical complement activation. Short-term treatment enhanced plaque and CAA binding of 3D6 with deposition of C3 fragments, red blood cell extravasation, and mild microhemorrhages. Repeated long-term immunization induced full complement activation, reduced total brain amyloid, and showed a trend toward vascular amyloid clearance, accompanied by BBB leakage and more microhemorrhages (ARIA-H). These effects were particularly evident in mice carrying human ApoE4, where the high CAA burden likely exacerbates complement activation and vascular injury, underscoring the clinical relevance of this model. While our findings establish complement activation as a mechanistic basis for ARIA, several limitations exist. First, we were unable to detect ARIA-E by MRI in mice, however the extravasation of FITC-Dextran and RBCs that were detected may be a close surrogate for ARIA-E. Second, we focused on the classical complement system due to its role in opsonization and phagocytic clearance of antibody-antigen complexes; it remains possible that the alternative and/or lectin complement pathways may also contribute to ARIA. Third, although we performed RNA sequencing on bulk brain tissue and isolated endothelial cells, we cannot define the gene expression changes in specific immune or vascular cells but plan to do so in future studies. Fourth, the mouse complement system is not identical to that in humans but complement appears to play a strong role in ARIA in both species. In summary, CAA-associated complement activation emerges as a key driver of anti-amyloid-associated ARIA, highlighting the potential of complement inhibition, possibly through blocking C1q binding to the anti-amyloid antibody or by modulating the proteolytic cleaves and/or regulatory proteins in the complement cascade to mitigate antibody-induced vascular injury while preserving therapeutic efficacy.

## Conclusion

In summary, our findings reveal a clear mechanistic link between anti-Aβ immunotherapy, complement activation at CAA-laden vessels, and downstream vascular injury. Early antibody engagement of vascular amyloid initiates classical complement activation, while prolonged treatment drives full complement cascade engagement, BBB disruption, and microhemorrhages. These effects coincide with the recruitment and activation of C3-associated CD206⁺ perivascular macrophages, extracellular matrix remodeling, and increased vascular permeability, collectively indicating that complement-mediated opsonization and inflammatory damage -- not simple Aβ redistribution -- underlie the observed cerebrovascular fragility. Although these results deepen our understanding of how complement-driven immune responses shape vascular outcomes during anti-Aβ immunotherapy, further work is needed to determine whether lowering complement activation, either genetically or pharmacologically, can reduce the risk of ARIA and mitigate the vascular vulnerability associated with anti-Aβ immunotherapy.

## Methods

### Animals

For the passive immunization experiments, different mouse models were used as described in **Table 1**. For the acute immunization studies, aged *APP^NL-G-F^* homozygous knock-in mice (abbreviated as *APP^NL-G-F^* KI mice in this study) were used. For the short and chronic anti-Aβ passive immunization studies, *APP/PS1dE9;APOE4* mice (*APPswe/PS1*dE9 mice crossed with human *APOE4* targeted-replacement mice; breeders were obtained from Dr. Martine Sadowski, NYU as previously described [33]. The experiments were conducted in aged mice (>16-mo-old) after they had developed CAA. Mouse genotypes were confirmed from a tail biopsy using Transnetyx automated genotyping services.

**Table 1:**
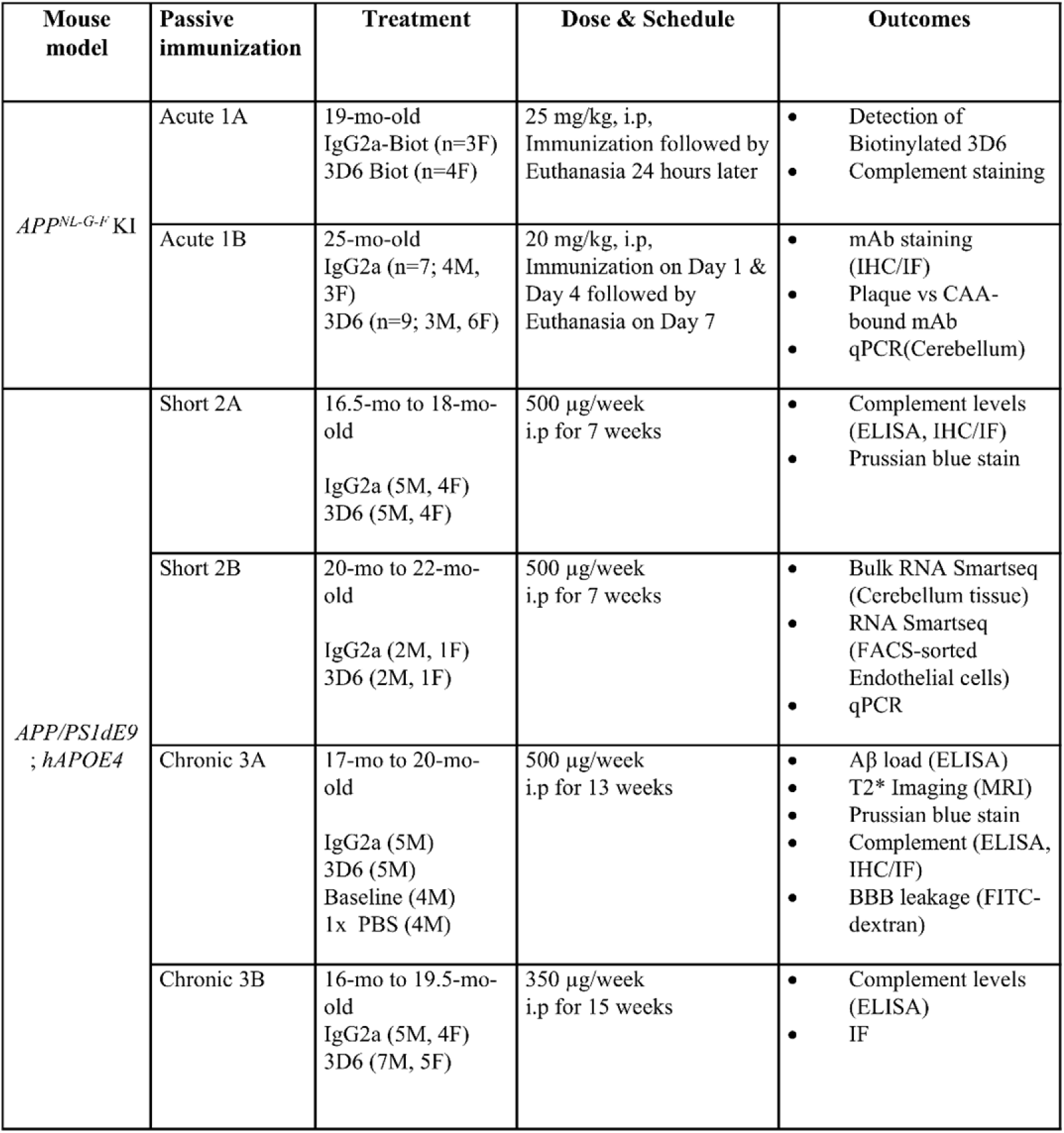
Anti-Aβ mAb passive immunization and mouse models.

Experimental procedures were carried out in compliance with institutional guidelines (IACUC, Institutional Animal Care and Use Committee) at Brigham and Women’s Hospital as per the Guide for the Care and Use of Laboratory Animals. Brigham and Women’s hospital’s assurance (A4752-1) is available on file with the Office of Laboratory Animal Welfare (OLAW). Mice were housed 2-5 per cage with a 12:12 h light/dark cycle with food and water provided *ad libitum*.

### 3D6 Antibody Generation

The original mouse antibody subtype of 3D6 is IgG2b [93], however we cloned it into an IgG2a subtype to obtain the mouse equivalent of the human IgG1 bapineuzumab antibody with analogous ADC effector functions. First, the variable regions of the light and heavy chains of the murine antibody 3D6 were amplified. In the next step, these were ligated into the dual expression vector pVITRO2-neo-mcs (InvivoGen Europe) upstream of the respective constant regions of the antibody, mouse kappa and mouse IgG2a. FreeStyle™ 293-F cells (Gibco) were then transfected with this vector for the constitutive co-expression of two genes. The cells were then converted into a stable cell line using G418 (Geneticin, InvivoGen) through the neomycin resistance of the vector. Freestyle 293-Expression Medium (Gibco) was inoculated with stable 3D6 IgG2a expressing HEK923F cells. Cells were cultivated at 37°C, 8 % CO2 and 3 mM Sodiumbutyrate at 125 rpm for three days. Cells were harvested and the supernatant was prepared for antibody purification. In preparation of affinity purification on protein G Sepharose, supernatant was mixed with binding buffer (200 mM Na2HPO4, 750 mM NaCl, pH 7.0) in a 5:1 ratio and applied to a 5 ml HiTrap Protein G HP column (Cytiva Europe GmbH). Unspecific bound proteins were removed by running a washing step with 40 mM Na2HPO4 buffer pH 7.0, containing 1M NaCl. The bound antibody was recovered by an acid elution step (Elution buffer: 0.1M glycine-HCl, pH 2.7) immediately followed by neutralization (Neutralizing buffer: 1 M Tris-HCl, pH 9.0). The antibody was dialyzed against PBS buffer (2,7 mM KCl, 1,7 mM KH2PO4, 137 mM NaCl, 8.1 mM Na2HPO4, pH 7.13). Subsequently, the dialyzed sample was sterile filtered and concentrated to 5.0 mg/ml by use of a VIVASPIN 20 concentrator (Satorius). Protein concentration was measured at 280 nm and computed by means of the molar extinction coefficient (Abs 0.1% (=1 g/l) 1.550).

### Anti-Aꞵ Antibody Treatments

Acute, short-term and chronic anti-Aβ passive immunization paradigms were used to evaluate the effect of anti-Aβ antibody-induced vascular side-effects known as ARIA. Details for each study are provided in **Table 1**. Recombinant 3D6 anti-Aβ antibody, the murine IgG2a precursor to bapineuzumab, and a murine IgG2a isotype control antibody were generated as described above by Drs. Stephan Schilling and Jens-Ulrich Rahfeld (Fraunhofer Institute for Cell Therapy and Immunology, Halle (Saale), Germany). We designed the following experiments to determine the temporal binding of 3D6 and its effects on the cerebrovasculature: 1) Acute studies to determine initial binding sites and complement activation in brain: 1A. Single intraperitoneal (i.p.) injection of 25 mg/kg 3D6-biotinylated or IgG2a-biotinylated antibodies (conjugated with biotin as per manufacturer instructions; Pierce™ Premium Grade Sulfo-NHS-LC-Biotin #PG82075, Thermo Scientific™) in 19-mo-old *APP^NL-G-F^* KI mice followed by euthanasia and brain harvest 24 hours later; 1B. Two-dose i.p. immunization with 20 mg/kg 3D6 or IgG2a on Days 1 and 4 followed by euthanasia on Day 7 in 25-mo-old *APP^NL-G-F^* KI; 2) Short-term studies to determine intermediate effects: 2A. 7-week treatment of weekly 500 µg 3D6 or IgG2a i.p immunization in 16.5-mo-old *APP/PS1dE9;hAPOE4* mice for pathological and biochemical analyses; 2B. 7-week treatment of weekly 500 µg 3D6 or IgG2a i.p. immunization in 20-mo-old *APP/PS1dE9;hAPOE4* mice for brain and endothelial transcriptomics; and 3) Chronic studies to examine complement and microhemorrhages: 3A. 13-week treatment of weekly 500 µg 3D6 or IgG2a i.p. immunization in 17-mo-old *APP/PS1dE9;hAPOE4* mice; 3B. 15 weekly treatments of 350 µg 3D6 or IgG2a i.p. immunization in 16-mo-old *APP/PS1dE9;hAPOE4* mice; 3C.13 weekly treatments of 500 µg 3D6 or IgG2a i.p. immunization in 22-mo-old *APP^NL-F^* KI mice. Mice were weight-balanced across groups in the short-term and chronic studies due to the uniform dose.

### Euthanasia and Tissue Collection

Mice were euthanized using CO2 inhalation and after bilateral thoracotomy, blood samples were drawn from the right cardiac ventricle followed by transcardial perfusion with ice cold 1x Phosphate buffered saline (PBS). Blood samples were later processed to separate serum and plasma with 50 mM EDTA. Brains were harvested and bisected sagittally. One hemibrain was snap-frozen in liquid nitrogen and stored at −80 °C until use. The other hemibrain was fixed overnight in 4 % paraformaldehyde (PFA) and later transferred through 10% and 30% sucrose gradients, followed by cryosectioning on a freezing microtome (Epredia HM 450 Sliding Microtome) at 30 µm thickness as previously described [33,94].

### Brain Homogenization and Protein Extraction

Brain tissue was dounce-homogenized with T-Per reagent (Thermo Scientific, USA #78510) with protease inhibitor cocktail set I (Calbiochem #539131) and phosphatase inhibitor cocktail set III (Calbiochem #524627) at 3000 rpm (25 strokes). Samples were sonicated and centrifuged at 100,000g at 4°C for 1 hr in a Beckman ultracentrifuge. Supernatants (Tper-soluble extracts) were aliquoted and stored at −80 °C until use. Insoluble pellets were resuspended in an ice-cold guanidine buffer (5M Guanidine HCL pH 8.0), dounce-homogenized as before and centrifuged again at 100,000g at 4°C for 1 hr. Supernatants (GHCl-soluble extracts) were aliquoted and stored at −80 °C until use. For normalization of ELISA data, protein concentrations of each sample were estimated using the Pierce BCA kit (Thermo Scientific, #23227) as per manufacturer instructions.

### MSD V-PLEX Plus Aβ ELISA (4G8 - Aβ Triplex 38/40/42)

T-per soluble and GHCl brain homogenates from all samples were analyzed for Aβx−38, Aβx−40 and Aβx−42 with MSD plate (Meso Scale Diagnostics, Rockville, MD, USA # K15199G) precoated with capture antibodies. The assay was carried out according to the manufacturer’s instructions. Briefly, the MSD plate was incubated with diluent for 1 hour at room temperature, followed by washes and incubation with 25 µL of detection antibody and 25 µL of prepared samples (1:5000 dilution) at room temperature for 2 hours. Following washes, the plate with a read buffer was analyzed using the MSD Discovery Workbench analysis software.

### ApoE ELISA

The levels of human apoE (hapoE) were measured by sandwich ELISA as previously reported [95]. In brief, hapoE levels from the Tper and GHCl soluble brain homogenates were measured using HJ15.6 and HJ15.4b as capture and detection antibodies, respectively. Recombinant apoE4 (Leinco) was used as the standard for the human apoE ELISA.

### Complement ELISAs

Mouse complement C1q, C3 and C5b-9 levels were measured in brain and blood by ELISA. Complement initiating factor C1q was measured using sandwich ELISA. Briefly, a 96-well high-binding microplate (Greiner Bio-One 96-Well High binding Polystyrene Microplates, #655085) was coated with rabbit anti-C1q (Abcam ab182451, 1:500, in 1x PBS buffer) overnight at 4 °C. Following PBS washes, 4% BSA in PBS-Tween (0.05%) blocking solution was added and incubated at 37°C for 1 hour to block non-specific binding. After removal of blocking solution, 100 µl diluted samples (T-per soluble brain homogenates, 1:200 and corresponding serum samples 1:200 in 4% BSA) were added in duplicate to wells and incubated for 2 hours at room temperature. C1q protein (Complement Technology, #M099) was used as a standard protein. Pierce Antibody Biotinylation Kit (Thermo Scientific #90407) was used for C1q ab182451 antibody biotinylation. Bound C1q was detected after incubating the biotinylated rabbit anti-C1q (1:1000) for 1 hour at 37 °C, followed by incubation with HRP-Conjugated Streptavidin (1:10000, Thermo Scientific, #N100) for 30 min at 37 °C. Following three PBS-Tween washes, enzyme substrate TMB (Thermo Scientific, # 34028) was added and after noting color development, the reaction was stopped using stop solution (Thermo Scientific #N600) and the plate was read at 450 nm using a BioTek 800 TS absorbance reader.

Mouse C3 levels were quantified as previously reported[96] with minor modifications. In brief, a high-binding microplate was coated with goat anti-mouse C3 antibody (1:500, 8 μg / mL, MP Biomedicals #55463) in 1x PBS overnight at 4 °C. Following three PBS-Tween washes, wells were blocked with 4% BSA-Tween 20 (0.05%) at 37°C for 1 hour.

After removing the blocking solution, diluted samples (T-per brain homogenates at 1:10; serum at 1:50,000 or depending on the experiment, plasma at 1:25,000 dilution with BSA) and standard C3 protein (Complement Technology, #M113) were added and incubated at room temperature for 2 hours. After three PBS-Tween washes, biotinylated goat anti-mouse C3 antibody (1:1000, MP Biomedicals #55463 biotinylated as described above for anti-C1q) was added and incubated for 1 hour at 37°C. Following washes, HRP-conjugated streptavidin was added and after three 1x PBS-Tween washes, enzyme substrate TMB (Thermo Scientific, # 34028) was added. Finally, upon color development, stop solution was added and absorbance was read at 450 nm as mentioned above.

Mouse terminal complement complex, C5b-9 was assayed using the Sandwich-ELISA Agilent AFG Scientific Mouse Terminal Complement Complex C5b-9, TCC C5b-9 ELISA kit (#EK730329) as per manufacturer instructions. In brief, standards or samples (1:5 dilution) were added to the pre-coated plate containing antibody specific to the terminal complement complex C5b-9, TCC C5b-9. Later HRP-conjugated antibody for TCC complex was added, followed by addition of TMB substrate solution and stop solution. The optical density (OD) was measured at 450 nm wavelength. One sample each from IgG2a and 3D6 treated mice were removed as they were identified as outliers via ROUT analysis.

### Histology and Imaging

Immunofluorescence staining: Brain sagittal sections were cut on a freezing microtome at 30 µm thickness followed by storage in 1x PBS buffer with 0.02% sodium azide. Brain sections underwent either free floating or slide immunostaining. In brief, brain tissue sections were washed in 1× PBS for 3 times 5 min each. Following washes in PBS, sections were blocked either in 10% normal goat serum (Gibco, #16210064), or heat inactivated goat serum (Novus Biologicals #S13110H), or normal donkey serum (Sigma, #S30-M) for 1 hour at room temperature followed by overnight 4°C incubation with primary antibodies against C1q (1:500, Abcam #ab182451), C3b/iC3b/C3c (1:300, HycultBiotech #HM1078), C5b-9 (1:300, Bioss #bs-2673R), CD31 (1:500, R&D Systems #AF3628), C3 (1:200, HycultBiotech #HM1045), GFAP (1:1000, Abcam # ab5804), MMR/CD206 (1:300,R&D #AF2535) MMP-9 (1:500, Bioss #bs-0397R), Iba-1 (1:1000; Dako, Wako #019-19741), Iba-1 (1:500, Synaptic Systems, #234308), CD68 (1:500, Bio-Rad #MCA1957), S97 Aβ (pan-Aβ rabbit polyclonal, 1:2000, gift Dr. Dominic Walsh, BWH, USA), CD31 (1:500, R&D #AF3628), Collagen-IV (1:300, Abcam #6586), Alexa Fluor 488 anti-β-Amyloid, aggregated antibody (1: 200, clone A17171C, Biolegend # 856508), Von Willebrand Factor Antibody (1:500, Novus Biologicals #NB600-586). Sections were washed 3 times in 1x PBS, 5 min each followed by 2 hour incubation at room temperature with corresponding secondary antibodies Goat anti-rabbit IgG AF 555, Invitrogen #A-21430, Goat anti-mouse IgG2a AF 647, Invitrogen #A-21241, Goat anti-mouse IgG2a AF 488, Invitrogen #A-21131, Goat anti-rat IgG AF 555, Invitrogen #A-21434, Donkey Anti-Rabbit IgG AF 647, Invitrogen #A-31573, Donkey Anti-Goat IgG Cy3, Jackson ImmunoResearch #705-165-147, Donkey Anti-Rat IgG Cy3, Jackson ImmunoResearch #712-165-150, Donkey Anti-Goat IgG CF 750, Biotium #20362, Donkey Anti-Guinea Pig IgG AF 488, Jackson ImmunoResearch #706-545-148, Donkey Anti-Rabbit IgG AF 488, Jackson ImmunoResearch #711-545-152, Donkey Anti-Goat IgG AF 633, Invitrogen # A-21082, Donkey Anti-Rabbit IgG AF 647, Invitrogen #A32795. After 3 washes in 1x PBS for 5 min each, sections were stained for fibrillar amyloid by incubating in Amylo-Glo dye solution (1x, Biosensis # TR-300-AG) for 10 min as per manufacturer instructions and later transferred to Superfrost Plus glass slides and coverslipped with custom made PVA-DABCO aqueous mounting media.

Fluorescent staining was visualized using a Axioscan 7 KMAT Slide scanner, 20x objective, with 2 to 3 µm z-stacks or Nikon Ti Eclipse Spinning Disk confocal microscope, 40x objective, 1 µm z-stacks. Areas of interest including cortex, hippocampus and cerebellum and images were taken from 4–5 sagittal tissue sections per mouse. Images were analyzed for % area fractions and colocalization data using custom made Fiji ImageJ macros [97] for % area fraction or object based image colocalization analysis between markers (supporting material). For CAA-associated complement analysis, individual CAA vessels (on average 22 to 25 vessels/mice) were outlined.

To detect microhemorrhages along with vascular amyloid and complement deposition, double IHC staining was performed for complement iC3b and S97 Aβ. Briefly, brain sections were mounted on the glass slides, air dried and rehydrated. After incubation in 1x PBS, tissue sections were treated with BLOXALL Endogenous Blocking Solution for Peroxidase and Alkaline Phosphatase (Vector Laboratories, #SP-6000-100) for 10 min. Following washes, sections were blocked with SuperBlock Blocking Buffer (ThermoFischer, # 37536) for 1 hour at room temperature. Sections were then incubated overnight at 4°C with primary antibody against C3b/iC3b/C3c (1:300, HycultBiotech #HM1078). The next day after 1x PBS washes, sections were incubated with secondary antibody, biotinylated Goat Anti-Rat IgG Antibody, Mouse Adsorbed (1:500, Vector Laboratories, #BA9401.5) followed by 1x PBS washes. Vectastain ABC kit (Vector Laboratories, #PK-6100) was used to label biotin with HRP as per the manufacturer’s instructions and visualized with a 3,3′-Diaminobenzidine (DAB) staining kit (Vector Laboratories, #SK-4100). Following Prussian blue staining as discussed below, sections were blocked with SuperBlock Blocking Buffer and overnight 4°C incubation with primary antibody against S97 Aβ (1:2000), washes with 1x PBS and 30 min secondary antibody incubation with Goat Anti-Rabbit IgG conjugated with Alkaline Phosphatase (1:100, SouthernBiotech, #4050-04), visualized with Vector Red substrate kit (Vector Laboratories, #SK-5100). After dehydration and clearing tissues with Histoclear, slides were coverslipped with DPX mounting media and imaged with 20x objective on Axioscan 7 KMAT Slide scanner and Nikon Eclipse E400 microscope.

### Microhemorrhage Detection

Prussian blue (PB) staining was performed as previously described [94]. Briefly, 4-5 sagittal mouse brain sections were mounted on glass slides, air dried, followed by rehydration in MilliQ water and 30 min incubation in 2 % potassium hexacyanoferrate trihydrate (Sigma, St. Louis, MO, USA; P3289) dissolved in 2 % Hydrochloric acid. After a quick rinse in MilliQ water, sections were counterstained with Nuclear Fast Red Counterstain (Vector Laboratories, Inc, H-3403-500) followed by dehydration steps in ethanol and Histoclear. Coverslip was done in DPX mounting medium and imaged on Axioscan 7 KMAT Slide scanner.

### Hematoxylin & Eosin Stain (H&E)

Hematoxylin & Eosin stain was performed as per the manufacturer instructions (VB-3000, VitroView Hematoxylin and Eosin Stain Kit). In some stainings to detect hemorrhages and red blood cell extravasation, prussian blue stain was followed by Eosin only counterstain. Eosin was shown to detect red blood cytoplasm as orange-red color [98].

### Magnetic Resonance Imaging (MRI)

In vivo brain imaging was performed in 20-month-old *APP/PS1dE9;hAPOE4* mice that were chronically treated with 500 µg/week for 13 weeks either with 3D6 mAb or 1x PBS. Imaging was done using a 7.0T Bruker BioSpec® USR magnet equipped with cryoprobe technology for high spatial resolution imaging of small models. Images were taken using T2 Turbo RARE_3D_Inv_3D (FLAIR), SWI_FcFLASH_3D and T2*-FLASH weighted gradient echo to detect edema and hemosiderin deposits resulting from cerebral microbleeds (cMBs). The voxels are roughly 0.07x0.07x0.7mm with a 0.3mm slice gap. We encountered artifacts along the pial surface air/brain border using the SWI and FLAIR sequences, however, we were able to get detectable hemorrhages using the T2*-FLASH weighted sequence. Nineteen slices covering the whole brain were analyzed. Hypointense regions in T2*-weighted images considered to be cMBs were verified by comparison to Prussian blue histology staining in the similar focal plane and region of interest in the whole brain. As a control group for detecting spontaneous hemorrhages, 1x PBS treated mice were used.

### BBB Permeability Assessment

After the last MRI scan, BBB leakage or disruption was visualized using dextran conjugated fluorophore using the modified protocol[99,100]. Briefly, mice received intraperitoneal injection of Anionic, Lysine Fixable, Dextran-fluorescein, 3 kDa (10mg/kg body weight, 10 mg/ml in saline, Invitrogen #D3306) approximately 3.5 hours before euthanasia. After perfusion with 1x PBS, brains were harvested and processed as above. Slides Scanned images were analyzed for dextran extravasation with z-stacks of 3 μm plane intervals and a resolution of 0.325 μm per pixel. ImageJ macro was used to evaluate the % Dextran area.

### RNA Sequencing

Changes in gene expression were determined by RNA extraction from the cerebellum, due to its propensity for CAA and microhemorrhages, and FACS sorted cerebrovascular endothelial cells and sequenced using Smart-seq2. Briefly, after immunization, the brain was harvested and a small portion of cerebellum was snap-frozen, while the rest of the brain (remaining tissue without olfactory bulb) was used to FACS-sort endothelial cells as per established protocols [101–103] with some modifications. Cerebellar tissue was lysed with a pestle and RNeasy Plus Mini Kit (Qiagen #74134) was used for RNA extraction. RNA Integrity Number (RIN) was measured with High Sensitivity Agilent Tape Station (Genomics and Technology Core Facility, MGH). Samples with RIN >7.3 were used for the sequencing. For endothelial cell isolation, the left-over hemi-brain tissue including cerebellum was digested with collagenase-D (2 mg/ml, Roche #11088858001) in HBSS buffer (Gibco, #14175095) at 37°C for 1 hour at 500 rpm, followed by a secondary digestion with collagenase/dispase (2 mg/ml, Roche #11097113001) in 2 % fetal bovine serum (FBS) and deoxyribonuclease I (25µg/ml, Worthington # LS002139) at 37 °C for 40 min at 500 rpm. Myelin removal was achieved by adding 25% bovine serum albumin (BSA) and centrifuging at 1000 g for 30 min at 4°C. The pellet was washed in 2% FBS in 1x PBS and then stained with CD13 FITC (BD Pharmingen #558744), CD31 APC (BD Pharmingen #551262), CD41 PE (BD Pharmingen #558040), and CD45 (BD Pharmingen #553081) for 20 min at 4°C. Cells were later stained with DAPI (100 ng/ml, BD Pharmingen #564907) for 5 min before washing with 2% FBS in 1x PBS. Following centrifugation, the sample was filtered through a 40 µm FACS tube and analyzed on BD FACS Standard Aria IIu. CD31^+^CD13^-^CD41^-^CD45^-^DAPI^-^ endothelial cells (average 20,000) were FACS-sorted and RNA was extracted using an Arcturus PicoPure RNA Isolation Kit as per manufacturer instructions (Applied Biosystems #12204-01). RNA integrity was analyzed using the Tape Station and 0.5 ng/ul RNA was sent to Broad Technology Labs and sequenced by the Broad Genomics Platform using the Smartseq2 protocol. RNA extracted from the right cerebellar hemisphere was analyzed by bulk RNA sequencing using Smart-seq2 (2×38 bp paired-end), yielding a median of 9.3 million reads, 94.0% alignment, and >10⁴ detected genes per sample. RNA-seq analysis was performed on Linux distribution. Quality control data was generated from the Fastq files using the FastQC tool; any adapter sequences and low-quality base ends were trimmed using TrimeGalore, trimmed reads are mapped to the GRCm39 build of the Mus musculus reference genome using STAR (Spliced Transcripts Alignment to a Reference) aligner. Duplicate reads were removed with Picard, followed by generating BAM files. Later, FeatureCounts were used to extract the gene count data. Raw counts were normalized using DESeq2 (v1.30.1) on “R”. Hypothesis testing was performed using DESeq2’s Wald test. Adjusted p-values were determined using the Benjamini-Hochberg procedure. Volcano plots were generated in “R” using the EnhancedVolcano (v1.8) package with fold change cutoffs >1 and adjusted p-value <0.05. Heatmaps and bar-plots with pathways were generated through “pheatmap” and Enrichplot packages respectively.

### qPCR

For qRT-PCR analysis, RNA was extracted from the cerebellum tissue and concentration measured on Nanodrop (NanoDrop ND-1000 Spectrophotometer). Single-stranded cDNA was synthesized from 100 ng of the total RNA using a High-Capacity cDNA Reverse Transcription kit (Applied Biosystems #4368814) following the manufacturer instructions. The amplification reactions were carried out in the Real-Time PCR System (Applied Biosystems) with the SYBR® Green Master Mix (Applied Biosystems #4309155). Technical duplicates of each biological sample were performed. β-actin was used as a housekeeping reference gene and relative mRNA expression Log 2^-(DDCT) between groups was estimated. The primer sequences used are listed in **Table 2**.

**Table 2:**
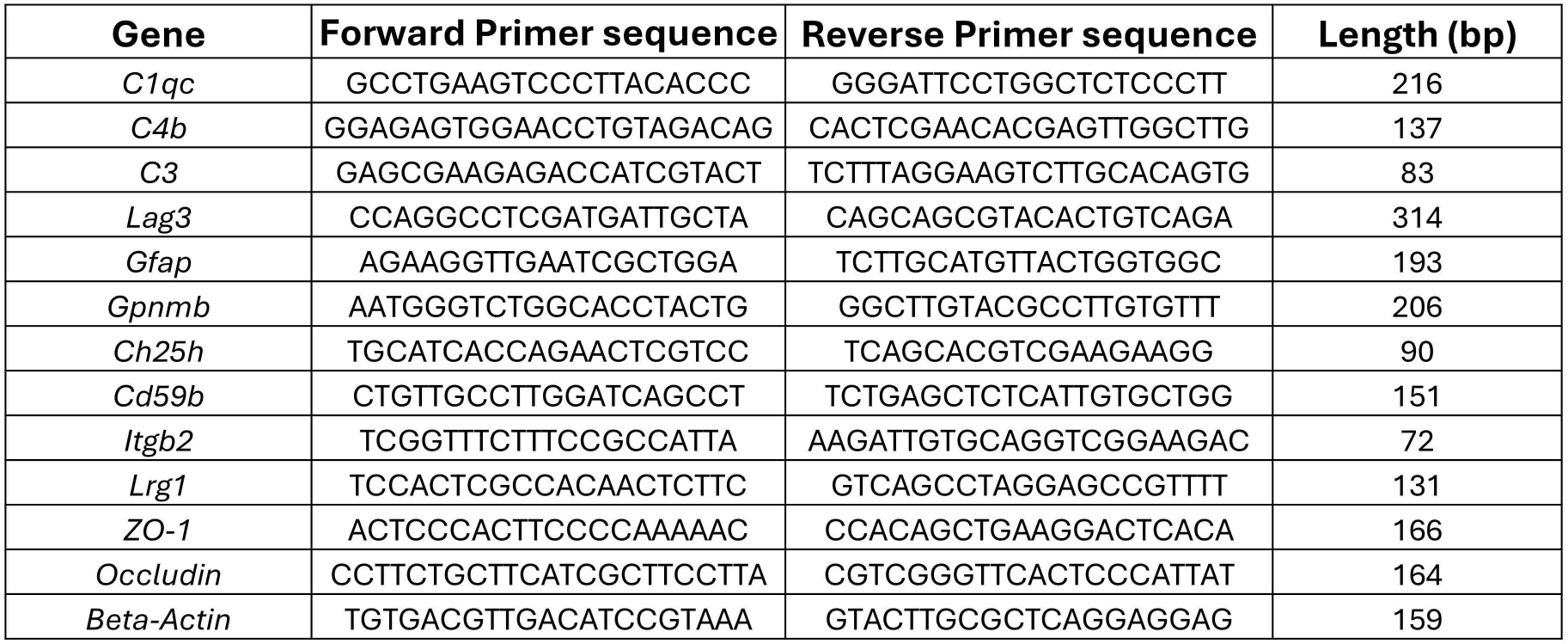
Primer list.

### Statistical Analyses

Statistics were performed using Graphpad Prism 10.0 (San Diego, USA). Normality of the residuals was analyzed using Anderson-Darling (A2*) test and differences in standard deviation through the Brown Forsythe test. Unpaired t test was used for Abeta ELISA. Normally distributed data was analyzed using parametric student’s-t-test and non-parametric data was analyzed either with Kruskal-Wallis or Manny-Whitney test. One-way ANOVA was used to compare all the groups in some cases. When there were no sex differences observed, data was pooled together to perform normal t-test between the groups. Data were expressed as mean ± SEM. Differences among groups were considered significant at values of p < 0.05. Outliers were removed via ROUT analysis. One outlier in each group was removed in the C5b-9 serum ELISA. For RNAseq data analysis “R” program (R 4.3.2) was used to generate the differential gene expression data. For a corrected measure of statistical significance that accounts for multiple testing, Benjamini-Hochberg procedure (FDR) was used and data was considered significant with adjusted p-value <0.05. Pairwise comparisons were made for mAb binding to plaques and vascular amyloid.

## Supporting information

Bathini et al. ImageJ macro script

Bathini et al. Complement ARIA_RNAseq supplementary document

## Data availability

Bulk RNA sequencing data and image analysis scripts are provided in the Supplementary Materials. All other data are available from the corresponding author upon reasonable request.

## Acknowledgments

We thank Maren Schroeder, Khyrul Khan, Jason Ciola, Brianna Rose Colletti, and Ilina Navani for their assistance with our mouse colony, and Martine Grenon and Maria-Tzousi Papavergi for their help with mouse brain tissue processing. We thank Dr. Michael C. Carroll for the C3 ELISA protocol and Dr. David Holtzman for the ApoE ELISA protocol used in our studies. We thank the NeuroTechnology Studio at Brigham and Women’s Hospital for providing access to the Axioscan 7 Slidescanner and the ARCND Flow Cytometry Core Facility for FACS sorting.

## Funding

This work was supported by NIH grants 1RF1 AG058657 and 1R01NS136122 and the Cure Alzheimer’s Fund (C.A.L.), NIH grant AG078106 (D.M.H.), and a Research Fellowship from the Alzheimer’s Association 23AARF-1029815 (P.B.).

## Authors’ contributions

P.B. and C.A.L. designed and planned all studies. J-U.R. and S.S. generated the treatment mAbs. D.M.H. and T.C.S. provided the mouse models. P.B. performed the experiments and analyzed the data. P.B. and C.A.L. interpreted the data and prepared the manuscript. All authors reviewed and approved the manuscript.

## Ethics declaration

### Competing interests

The authors declare no competing interests

### Consent for publication

Not applicable

## Conflict of interest statement

C.A.L. serves as a consultant or scientific advisory board member for Acumen, ADvantage Therapeutics, Apellis, Cyclo Therapeutics, Eli Lilly, Merck, MindImmune, Novo Nordisk, Receptive Bio, Switch Therapeutics and Therini Bio. D.M.H. co-founded and is on the scientific advisory board of C2N Diagnostics. D.M.H. is on the scientific advisory boards of Denali, Genentech, and Switch and consults for Pfizer, Roche, and Novartis. J-U.R. and S.S. are co-founders of Periotrap Pharmaceuticals. S.S. serves as consultant to Vivoryon Therapeutics N.V. All other authors have no conflicts to report.

## Abbreviations

3D6: Murine IgG2a precursor to bapineuzumab
Aβ: Amyloid-β (protein)
Abca1: ATP-binding cassette transporter A1 (gene)
AD: Alzheimer’s disease
ANOVA: Analysis of variance
ApoE / APOE: Apolipoprotein E
ApoE ε4 / ε4: Apolipoprotein E ε4 allele
ARIA: Amyloid-Related Imaging Abnormalities
ARIA-E: ARIA-edema (vasogenic edema or effusion)
ARIA-H: ARIA-hemorrhage (microhemorrhages or superficial siderosis)
BBB: Blood-brain barrier
C1q (C1qa, C1qb, C1qc): Complement component 1q (and its specific polypeptide chains)
C1r: Complement component 1r (serine protease)
C1s: Complement component 1s (serine protease)
C3: Complement component 3
C3a: Complement component 3a (anaphylatoxin)
C3aR: Complement component 3a receptor
C3b / iC3b / C3c: Complement component 3b / inactivated C3b / C3c (cleavage fragments)
C4 / C4b: Complement component 4 / 4b
C4b2b: C3 convertase (classical pathway)
C5: Complement component 5
C5a: Complement component 5a (anaphylatoxin)
C5aR: Complement component 5a receptor
C5b: Complement component 5b
C5b-9: Terminal membrane attack complex (MAC)
CAA: Cerebral amyloid angiopathy
CAA-ri: Cerebral amyloid angiopathy-related inflammation
Cfh: Complement factor H
Ch25h: Cholesterol 25-hydroxylase (gene)
DAPI: 4’,6-diamidino-2-phenylindole (fluorescent DNA stain)
DEGs: Differentially expressed genes
DESeq2: Differential gene expression analysis tool
df: Degrees of freedom
FACS: Fluorescence-activated cell sorting
Fc: Fragment crystallizable region (of an antibody)
FLAIR: Fluid-attenuated inversion recovery (MRI sequence)
Gfap: Glial fibrillary acidic protein (gene/marker)
Gpnmb: Glycoprotein nonmetastatic melanoma protein B (gene/marker)
H&E: Hematoxylin and eosin (stain)
i.p.: Intraperitoneally (injection method)
IgG2a: Immunoglobulin G subclass 2a
Lcn2: Lipocalin 2 (gene)
Lgals3: Galectin-3 (gene)
Lrg1: Leucine-rich alpha-2-glycoprotein 1 (gene)
Lyz2: Lysozyme 2 (gene)
mAbs: Monoclonal antibodies
MMP-9 / Mmp12: Matrix metallopeptidases
MRI: Magnetic resonance imaging
p / padj: P-value / adjusted p-value
PBS: Phosphate-buffered saline
PCA: Principal Component Analysis
qPCR: Quantitative polymerase chain reaction
RBC: Red blood cell
S97: A specific pan-Aβ antibody
t-PA: Tissue plasminogen activator

---

**Supplementary Fig.1:**
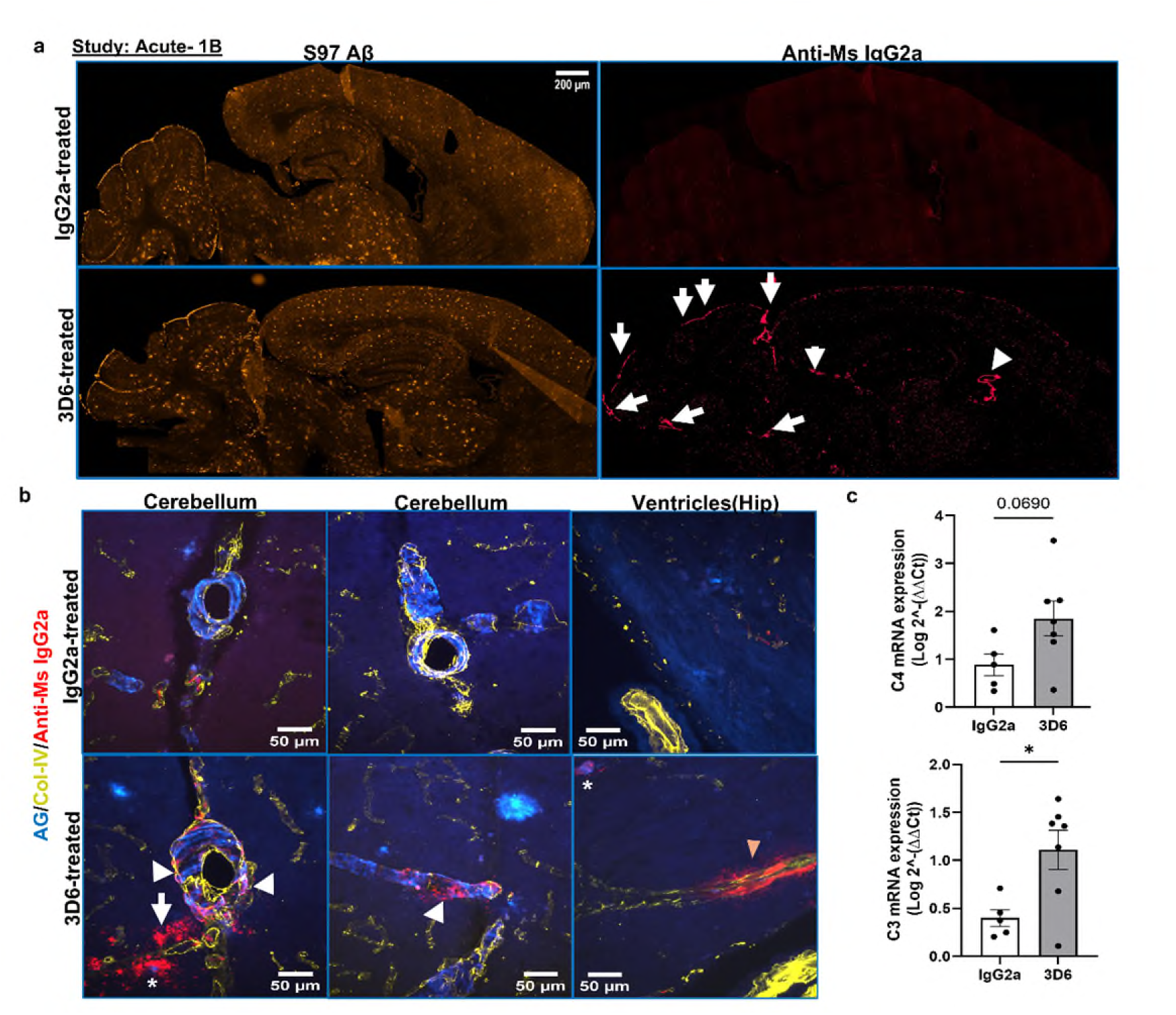
Acute anti-Aβ immunization indicates early preferential binding of 3D6 to vascular amyloid. **a,** Acute passive immunotherapy in 25-month-old *APP^NL-G-F^* KI mice with 20mg/kg 3D6 mAb on Day 1 and 4, followed by euthanasia on Day 7. Controls received IgG2a isotype control mAb. Immunostaining for S97 (general Aβ) and anti Ms-IgG2a (to detect 3D6 mAb) shows that 3D6 preferentially binds to amyloid (arrows) in the cortical and cerebellar meninges and large vessels near the ventricles. 3D6 can be seen in the choroid plexus as well (arrowhead). **b,** Brain sections stained with Amylo-Glo, Collagen-IV and anti-Mouse IgG2a. Cerebellar CAA-associated mAb binding is indicated by arrowheads, plaque labelling by asterisks, and non-CAA-associated vascular mAb near ventricles (right image panel) is indicated by a colored arrowhead. Unbound mAb in the parenchyma outside CAA vessels is indicated by an arrow**. c,** qPCR analysis for complement genes *C4* & *C3* from the bulk cerebellum tissue, Mann-Whitney test, *p < 0.05, data are expressed as mean ± SEM. Scale bar in a =200 µm, b = 50 µm.

**Supplementary Fig.2:**
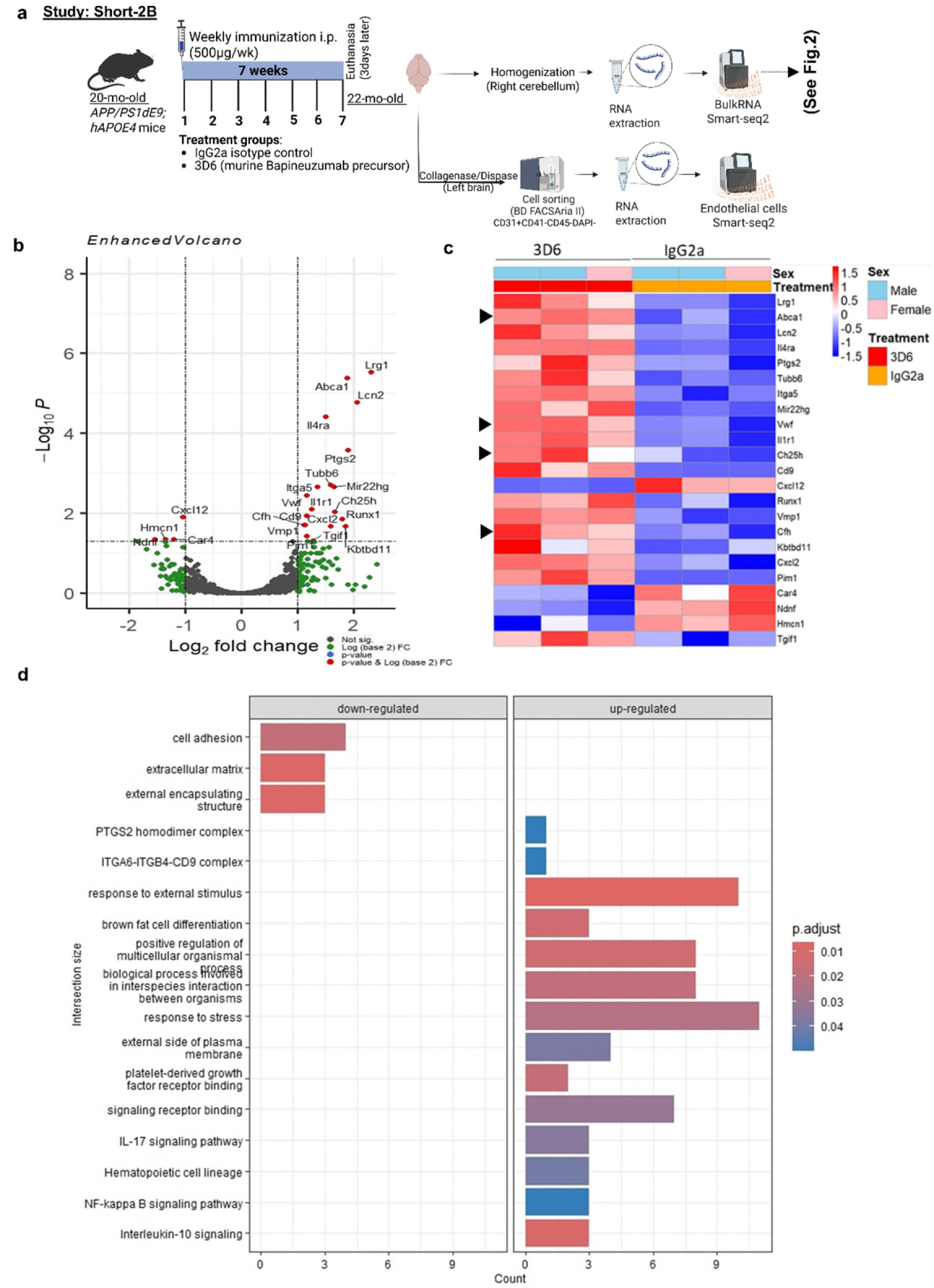
Passive anti-Aβ immunization for 7 weeks was associated with differentially expressed genes in the FACS-sorted CD31^+^ endothelial cells in the brain. **a,** Schematic for anti-Aβ immunotherapy in the 20-month-old *APP/PS1dE9;hAPOE4* mice with endothelial cell isolation, RNA extraction followed by Smart-seq2 (n=3, 2 Males and 1 Female mouse). **b,** Volcano plot showing differentially expressed genes. **c,** Pheatmap plot showing selected significantly expressed genes related to cholesterol efflux (*Abca1*, *Ch25h*) and blood homeostasis (*Vwf)* and complement genes *(Cfh)* and others related to lipid stress and inflammation, denoted with arrowheads. **d,** Bar plots representing the biological pathways that are down and up-regulated.

**Supplementary Fig.3:**
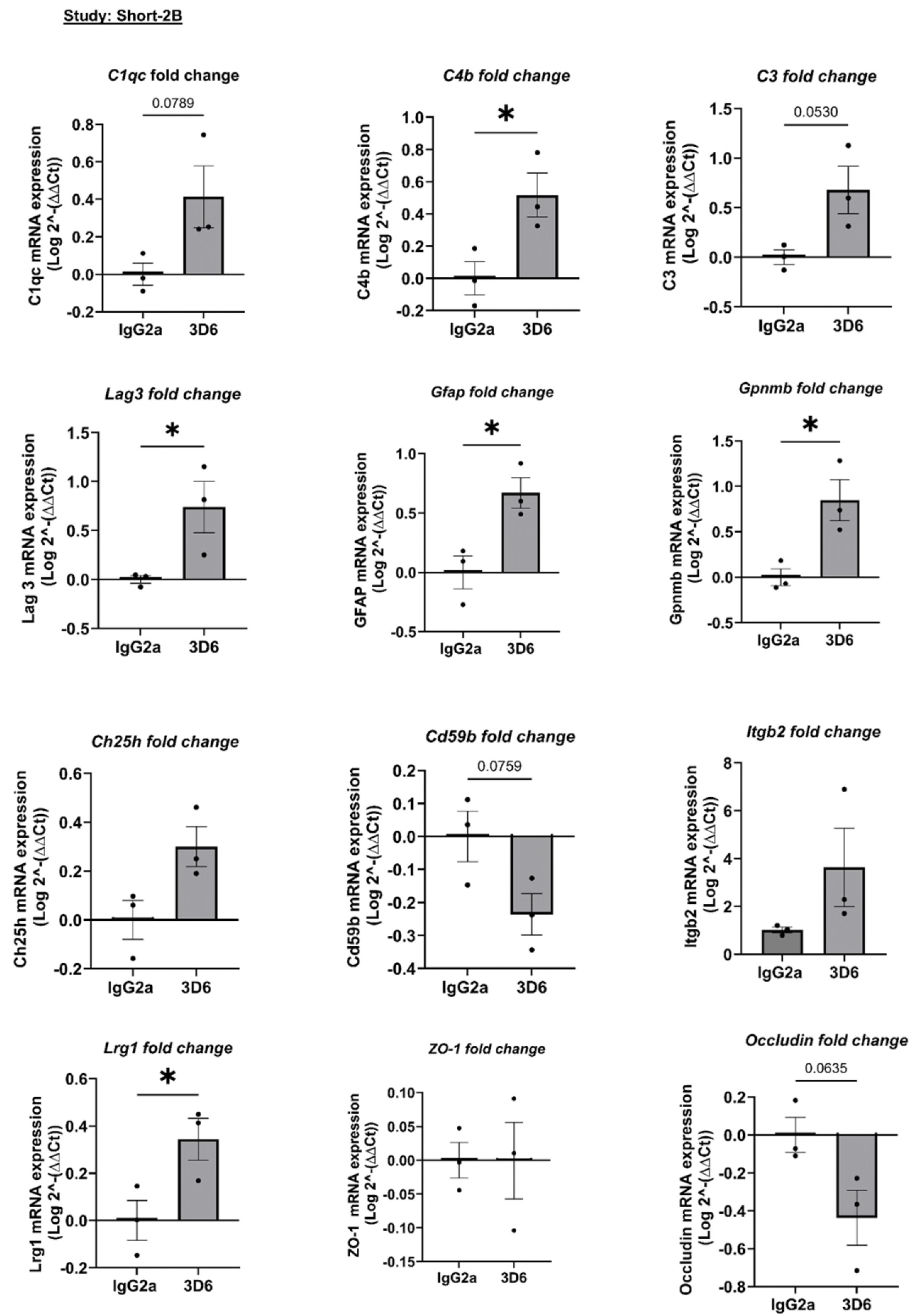
qPCR analysis of relevant genes associated with complement pathway and BBB integrity, following 7 weeks of passive anti-Aβ immunization in the 20-month-old *APP/PS1dE9;hAPOE4* mice. RNA was extracted, and qPCR was performed on the cerebellum tissue. Statistical analysis was conducted using a Parametric unpaired t-test *p<0.05. Data are expressed as mean ± SEM.

**Supplementary Fig.4:**
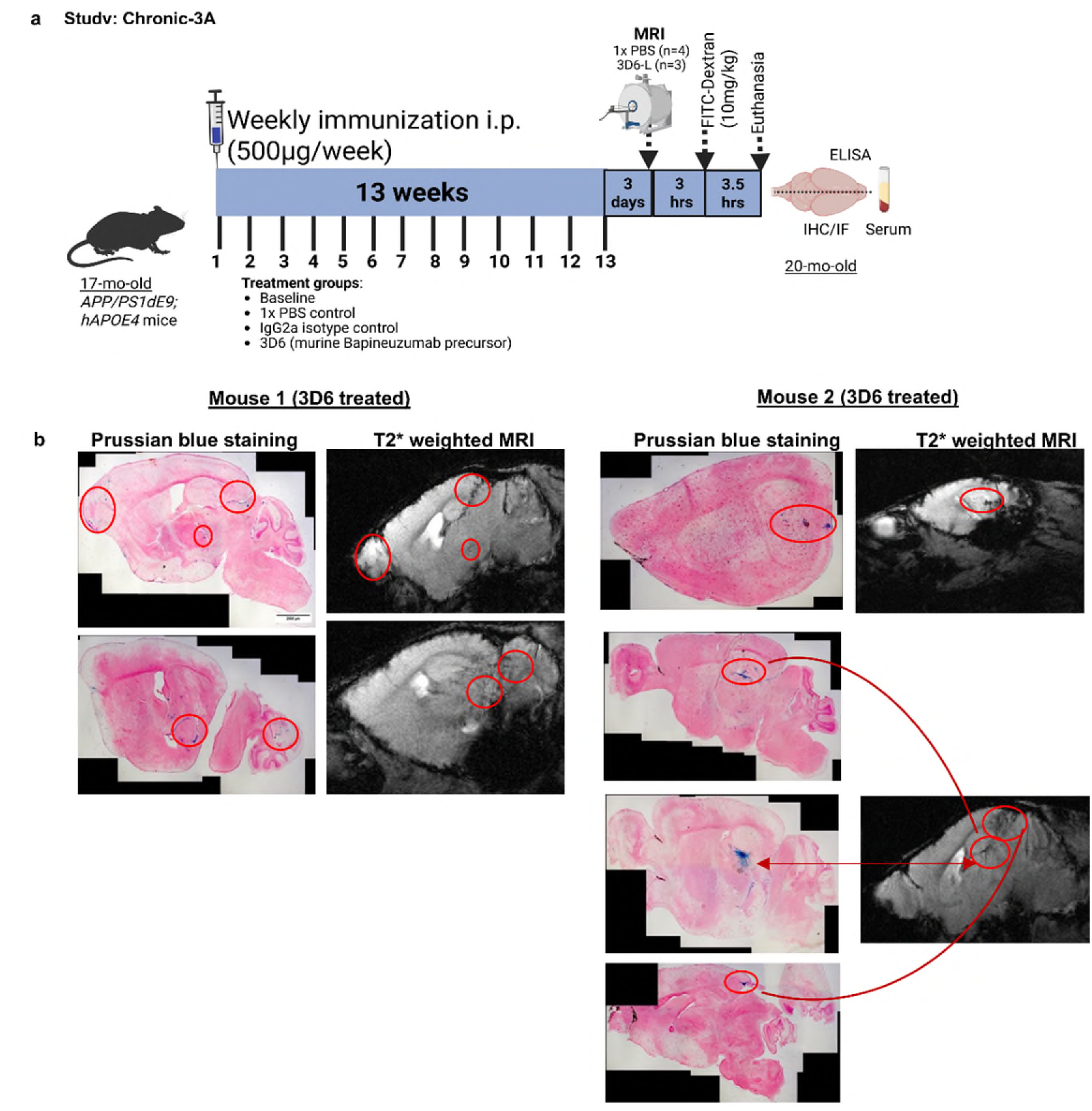
Anti-Aβ immunotherapy and associated microhemorrhages. **a,** MRI imaging was performed in 20-month-old *APP/PS1dE9;hApoE4* mice following 13 weeks of 500 µg/wk (15 mg/kg body weight) 3D6 mAb passive immunization and Prussian blue hemosiderin staining on the brain tissue sections after euthanasia. **b**, Representative Prussian blue stained brain sections from two 3D6 treated mice and their corresponding MRI sagittal planes (approximate) showing hemosiderin deposits and matched hypointense spots as indicated by circles or arrows.

**Supplementary Fig.5:**
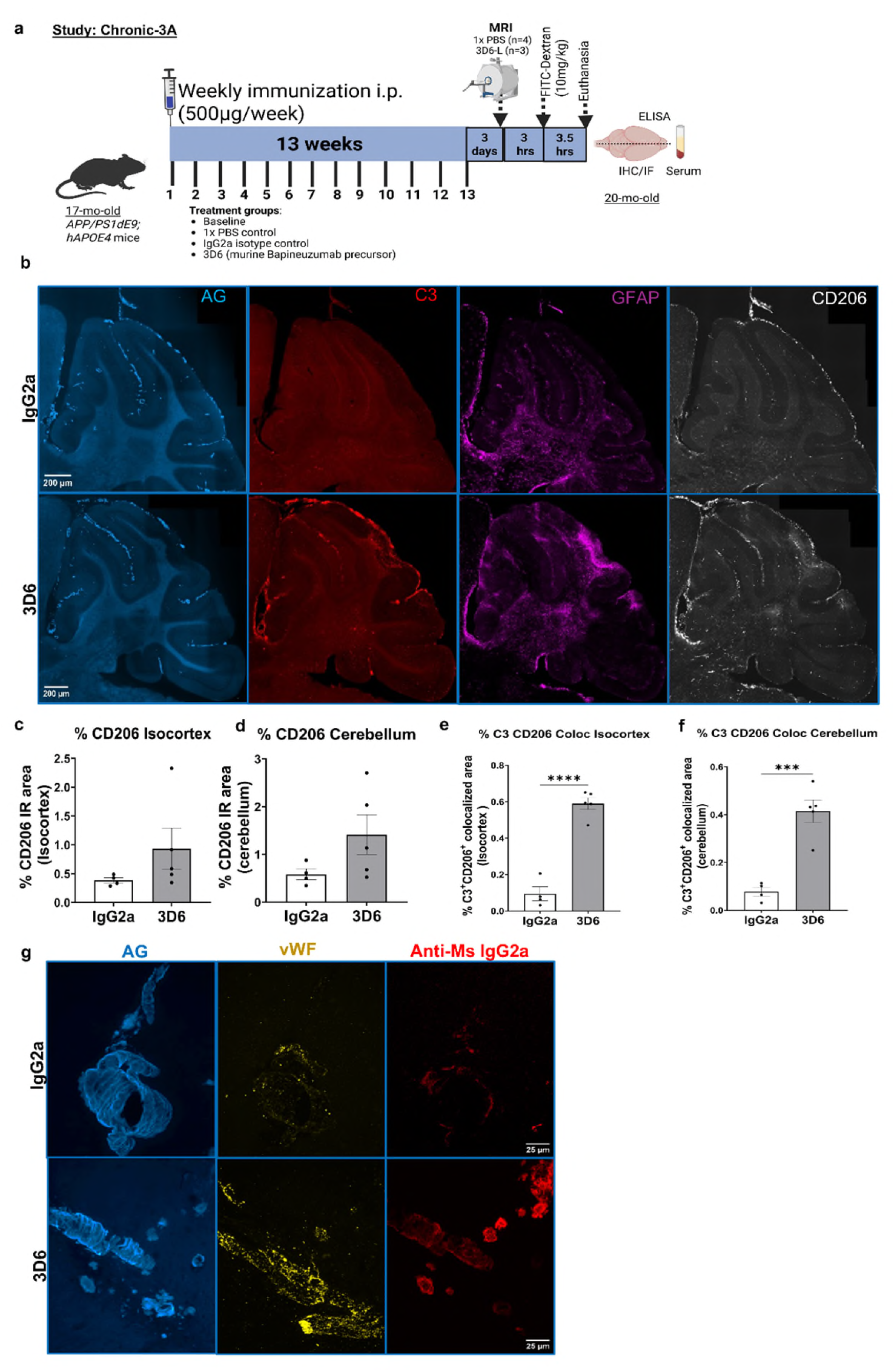
Chronic anti-Aβ passive immunization with 3D6 mAb caused increased recruitment of CD206+ border-associated macrophages (BAM) along C3 deposited areas and mAb-associated vWF deposition in aged 20-mo-old *APP/PS1dE9:hAPOE4* mice following 13 weeks of passive immunization with 500 µg/wk mAb. **a,** Schematic of passive immunization **b,** Representative Immunofluorescence staining in the cerebellum area for C3, GFAP, CD206, and Amylo-Glo (AG). ImageJ quantification of % CD206+ area in the **c,** Isocortex (cortex & hippocampus) and **d,** Cerebellum area. C3+CD206+ colocalized area in the **e,** Isocortex and **f,** Cerebellum area in IgG2a and 3D6 treated mice (n=4-5/group). Data represented as mean ± SEM. Parametric test with unpaired t-test ***p=0.0005 and ****p<0.0001. **g**, Immunostaining of brain sections with von Willebrand factor (vWF), a glycoprotein produced by endothelial cells or platelets, anti-mouse IgG2a, and amyloid staining with Amylo-Glo. Scale bar: b=200 µm, g= 25 µm.

## Notes

### Competing Interest Statement

The authors have declared no competing interest.

