## Supplementary material for "Early Binding of Anti-Amyloid Antibodies to CAA Drives Complement Activation, Inflammation and ARIA in Mice": Bathini et al. ImageJ macro script

### Hemosiderin (Prussian blue) analysis:

```
title = getTitle();

setTool("wand");

waitForUser("Draw an area of Interest");

roiManager("Add");

//10X Scaling slide scanner

run("Set Scale...", "distance=1 known=0.172 unit=um global");

run("Colour Deconvolution", "vectors=[H PAS]");

close();

selectImage (title + "-(Colour_1)");

run("8-bit");

setAutoThreshold("MaxEntropy no-reset");

//setThreshold(0, 202);

setOption("BlackBackground", false);

run("Convert to Mask");

waitForUser("erase the false staining");

selectImage (title + "-(Colour_1)");

roiManager("Select", 0);

roiManager("Set Color", "red");

roiManager("Set Line Width", 1);

run("Set Measurements...", "area mean integrated area_fraction display redirect=None
decimal=3");

run("Analyze Particles...", "size=5-Infinity show=Outlines display summarize add");

close("*");
```

### Object based co-localization (example):

```
// Choose the output folder
dir2=getDirectory("User Choose Result Data Folder");
originalName = getTitle();
title1=originalName;
//title1 = getTitle();
//title1=File.nameWithoutExtension;
//print(title1);


// 20x scaling slide scanner
run("Set Scale...", "distance=1 known=0.325 unit=um global");
run("Z Project...", "projection=[Max Intensity]");
run("Brightness/Contrast...");
run("Enhance Contrast", "saturated=0.35");
run("Stack to Images");

//ids=newArray(nImages);
ids=newArray(nImages);
for (i=0;i<nImages;i++) {
	selectImage(i+1);
	title = getTitle;
	print(title);
	ids[i]=getImageID;
	saveAs("Tiff", dir2+title); }

waitForUser(""); // to cross check everthing is in right direction


run("Images to Stack", "use keep");
roiManager("Show All");
//setTool("polygon");
run("Brightness/Contrast...");
waitForUser("Draw a region over the area of interest");
roiManager("Add");
//roiManager("Select", 0);
//run("Crop");
run("Deinterleave", "how=3 keep");



// Choose blue channel AmyloGlo
selectWindow("Stack #1");rename("AmyloGlo mask");
```

```
run("Subtract Background...", "rolling=200 sliding");
roiManager("Show All");
run("Set Measurements...", "area mean display redirect=None decimal=3");
roiManager("Select", 0);
run("Measure");
run("8-bit");
//run("Threshold...");
setAutoThreshold("Triangle dark");
//setThreshold(3, 255);
setOption("BlackBackground", false);
run("Convert to Mask");
waitForUser("");// to wait and confirm everything looks good



// Choose Yellow channel CD31
selectWindow("Stack #2");rename("CD31 mask");
run("Subtract Background...", "rolling=200 sliding");
roiManager("Show All");
run("Set Measurements...", "area mean display redirect=None decimal=3");
roiManager("Select", 0);
run("Measure");
run("8-bit");
run("Threshold...");
waitForUser ("Adjust the threshold then press Ok");



// Choose FR channel C1q
selectWindow("Stack #3");rename("C1q mask");
run("Subtract Background...", "rolling=200 sliding");
roiManager("Show All");
run("Set Measurements...", "area mean display redirect=None decimal=3");
roiManager("Select", 0);
run("Measure");
run("8-bit");
setAutoThreshold("Triangle dark");
//setThreshold(3, 255);
setOption("BlackBackground", false);
run("Convert to Mask");
waitForUser("");// to wait and confirm everything looks good
```

```
////////////// Object based colocalized spots AmyloGlo and C1q
imageCalculator("AND create", "AmyloGlo mask","C1q mask");
selectWindow("Result of AmyloGlo mask");rename("Result of AmyloGlo+C1q mask");
run("8-bit");
setAutoThreshold("Triangle dark");
//setThreshold(62, 255);
run("Invert");
run("Convert to Mask");


////////////// Object based colocalized spots CD31 and C1q
imageCalculator("AND create", "CD31 mask","C1q mask");
selectWindow("Result of CD31 mask");rename("Result of CD31+C1q mask");
run("8-bit");
setAutoThreshold("Triangle dark");
//setThreshold(62, 255);
run("Invert");
run("Convert to Mask");





// save mask file and analyze data

selectWindow("AmyloGlo mask"); roiManager("Show All without labels");
saveAs("Jpeg",dir2+title1+"_AmyloGlo mask.jpg");
RoiManager.setPosition(0);
roiManager("Select", 0);
run("Analyze Particles...", "size=0.5-Infinity show=[Overlay Masks] summarize");
//roiManager("Save", dir2 + title1 + AmyloGlomask + "RoiSet.zip");
run("Flatten");
saveAs("Jpeg",dir2+title1 +"_AmyloGlo mask.jpg");



selectWindow("CD31 mask");roiManager("Show All without labels");
saveAs("Jpeg",dir2+title1+"_CD31 mask.jpg");
RoiManager.setPosition(0);
 roiManager("Select", 0);
 run("Analyze Particles...", "size=0.5-Infinity show=[Overlay Masks] summarize");
```

```
 run("Flatten");
 saveAs("Jpeg",dir2+title1+"_CD31 mask.jpg");


selectImage("C1q mask");roiManager("Show All without labels");
saveAs("Jpeg",dir2+title1+"_C1q mask.jpg");
RoiManager.setPosition(0);
 roiManager("Select", 0);
 run("Analyze Particles...", "size=0.5-Infinity show=[Overlay Masks] summarize");
  run("Flatten");
  title1=originalName;
 saveAs("Jpeg",dir2+title1+"_C1q mask.jpg");



selectWindow("Result of AmyloGlo+C1q mask");roiManager("Show All without labels");
saveAs("Jpeg", dir2+title1+"_Result of AmyloGlo C1q overlay.jpg");
RoiManager.setPosition(0);
 run("Analyze Particles...", "size=0.5-Infinity show=[Overlay Masks] summarize");
 run("Flatten");
 saveAs("Jpeg", dir2+title1+"_Result of AmyloGlo C1q overlay.jpg");



  selectWindow("Result of CD31+C1q mask");roiManager("Show All without labels");
  saveAs("Jpeg",dir2+title1+"_Result of CD31 C1q overlay.jpg");
RoiManager.setPosition(0);
 roiManager("Select", 0);
 run("Analyze Particles...", "size=0.5-Infinity show=[Overlay Masks] summarize");
 run("Flatten");
 saveAs("Jpeg",dir2+title1+"_Result of CD31 C1q overlay.jpg");
  run("Close");
close("*");
```
